# MYC drives aggressive prostate cancer by disrupting transcriptional pause release at androgen receptor targets

**DOI:** 10.1101/2021.04.23.441016

**Authors:** Xintao Qiu, Nadia Boufaied, Tarek Hallal, Avery Feit, Anna de Polo, Adrienne M. Luoma, Janie Larocque, Giorgia Zadra, Yingtian Xie, Shengqing Gu, Qin Tang, Yi Zhang, Sudeepa Syamala, Ji-Heui Seo, Connor Bell, Edward O’Connor, Yang Liu, Edward M. Schaeffer, R. Jeffrey Karnes, Sheila Weinmann, Elai Davicioni, Paloma Cejas, Leigh Ellis, Massimo Loda, Kai W. Wucherpfennig, Mark M. Pomerantz, Daniel E. Spratt, Eva Corey, Matthew L. Freedman, X. Shirley Liu, Myles Brown, Henry W. Long, David P. Labbé

## Abstract

c-MYC (MYC) is a major driver of prostate cancer tumorigenesis and progression. Although MYC is overexpressed in both early and metastatic disease and associated with poor survival, its impact on prostate transcriptional reprogramming remains elusive. We demonstrate that MYC overexpression significantly diminishes the androgen receptor (AR) transcriptional program (the set of genes directly targeted by the AR protein) in luminal prostate cells without altering AR expression. Importantly, analyses of clinical specimens revealed that concurrent low AR and high MYC transcriptional programs accelerate prostate cancer progression toward a metastatic, castration-resistant disease. Data integration of single-cell transcriptomics together with ChIP-seq revealed an increased RNA polymerase II (Pol II) promoter-proximal pausing at AR-dependent genes following MYC overexpression without an accompanying deactivation of AR-bound enhancers. Altogether, our findings suggest that MYC overexpression antagonizes the canonical AR transcriptional program and contributes to prostate tumor initiation and progression by disrupting transcriptional pause release at AR-regulated genes.

**STATEMENT OF SIGNIFICANCE:** AR and MYC are key to prostate cancer etiology but our current understanding of their interplay is scarce. Here we show that the oncogenic transcription factor MYC can pause the transcriptional program of the master transcription factor in prostate cancer, AR, while turning on its own, even more lethal program.

## INTRODUCTION

Prostate cancer is the most common non-cutaneous malignancy and a leading cause of cancer-related lethality in men ^1^. The androgen receptor (AR), a ligand-activated transcription factor, is central to the homeostasis of normal prostate epithelium ^2,3^. Importantly, since the discovery that prostate cancer is reliant on androgen signaling to thrive ^4,5^, targeting AR activity continues to be the main pillar of prostate cancer therapy ^6^.

Prostate cancer initiation and progression involves the corruption of the normal prostate cancer transcriptional network ^7^. Loss of the *NKX3-1* homeobox gene is a frequent and early event in prostate cancer etiology while the *TMPRSS2-ERG* gene fusion and *FOXA1* mutations both identify major molecular subtypes of the disease ^8,9^.

Overexpression of c-Myc (MYC), a master transcription factor and oncoprotein whose expression and function are tightly controlled under normal circumstances, is frequently observed in prostate cancer. Nuclear overexpression of MYC protein is an early event observed in luminal cells of prostate intraepithelial neoplasia (PIN) and is maintained in a large proportion of primary carcinomas and metastatic disease ^10^. Importantly, about 25% of familial risk of prostate cancer map to germline variation at chromosome 8q24 with mechanistic evidence tying this region to *MYC* regulation ^11-13^. Critically, MYC overexpression in normal luminal cells of murine prostate is sufficient to initiate prostate cancer ^14^, providing evidence that deregulation of MYC protein expression is a critical oncogenic event driving prostate cancer initiation.

Although AR and MYC are both central to prostate cancer etiology, our current understanding of the interplay between these two transcription factors is scarce. A recent study revealed that MYC overexpression antagonizes androgen-induced gene expression in an androgen-sensitive cell line representative of advanced prostate cancer ^15^. However, it remains unknown how increased MYC expression shapes the AR transcriptional program in normal luminal prostate cells as they transition to PIN and subsequently progress from a localized to a metastatic disease.

Here we model MYC-driven prostate cancer initiation *in vivo* and define the transcriptional rewiring occurring in luminal cells at a single-cell level. We demonstrate that MYC overexpression diminishes the canonical AR transcriptional program, alters the AR cistrome, and results in the establishment of a corrupted AR transcriptional program in a murine model. We determine that an active MYC transcriptional program and low AR activity identify prostate cancer patients predisposed to fail standard-of-care therapies and most likely to develop metastatic castration resistant prostate cancer (mCRPC). Accordingly, we found that high *MYC* mRNA expression in castration-resistant tumors is also associated with a weakened canonical AR transcriptional program and a repurposing of the AR cistrome. Critically, integration of transcriptomic and epigenomic data reveals that MYC overexpression does not lead to the deactivation of AR-bound enhancers but instead results in RNA polymerase II (Pol II) promoter-proximal pausing at AR-dependent genes. Altogether, our findings suggest that MYC overexpression contributes to tumor initiation and progression by disrupting the AR transcriptional program.

## RESULTS

### MYC induces a profound transcriptional reprogramming in murine prostate lobes

To examine the transcriptional reprogramming associated with MYC-driven prostate cancer initiation, we compared a 12-week-old mouse that overexpresses an ARR_2_Pb driven human *c-MYC* transgene (*MYC*) in the prostate epithelium to a wild-type (WT) littermate ^14^. At 12 weeks of age, MYC overexpression induces cellular epithelium transformation to PIN, a premalignant condition that often precedes the development of invasive adenocarcinoma in humans ^16^, with varying penetrance across prostate lobes. Notably, the murine anterior prostate (AP) remained mostly unaffected by MYC overexpression while PIN penetrance reached 83% and 97% in the dorsolateral prostate (DLP) and ventral prostate (VP), respectively ^17^. Transcriptional profiling of whole prostate lobes at a single-cell level revealed a strong overlap with the matched bulk gene expression profiling across lobes and genotypes (WT and MYC; **Figure 1A-B** and **Supplementary Figure S1A**). Comparison of gene expression levels quantified by single-cell RNA-seq (scRNA-seq; aggregate expression) or bulk RNA-seq revealed that scRNA-seq quantitatively recapitulates bulk gene expression (**Figure 1C** and **Supplementary Figure S1B**). Accordingly, with the exception of the AP, unsupervised clustering revealed a strong correlation between single-cell transcriptome and the matched bulk transcriptome (**Figure 1D**) and revealed that MYC induces a profound transcriptional reprogramming in both the DLP and VP lobes (**Figure 1E**).

**Figure 1:**
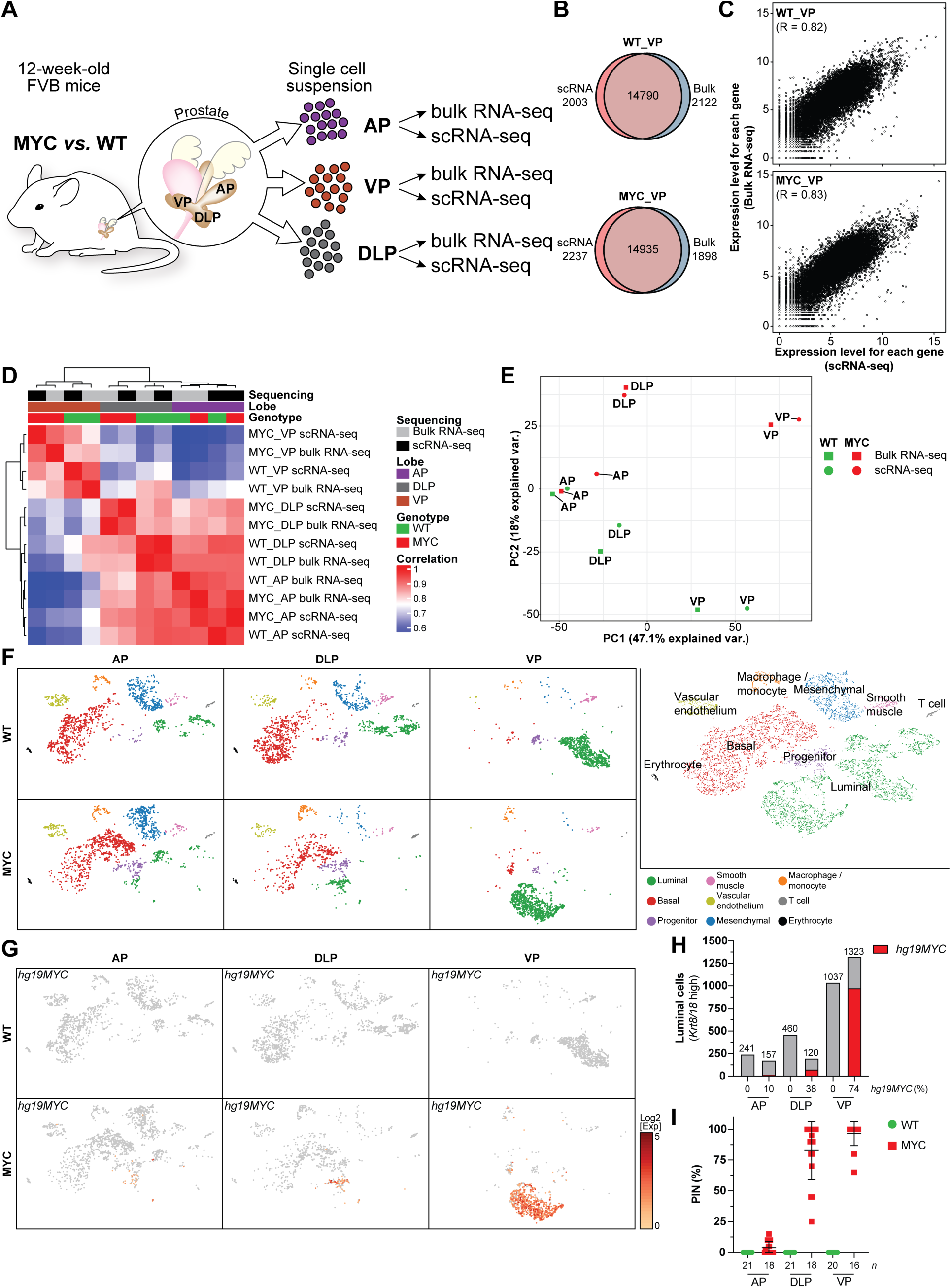
MYC induces a profound transcriptional reprogramming in murine prostate lobes. (**A**) Graphical summary of the experimental design. (**B, C**) Transcriptional profiling of WT and MYC-transformed VP reveal high concordance for the total number of genes detected (**B**) and their expression levels (**C**) between bulk and single-cell RNA-seq. (**D, E**) Sample-sample correlation (**D**) and principal component analysis (**E**) between bulk and matched single-cell transcriptome identifies distinct transcriptional profiles across murine prostate lobes. (**F**) Single-cell census of WT and MYC-transformed AP, DLP and VP. tSNE of scRNA-seq profiles colored using known markers identified nine major subpopulations across prostate lobes. (**G-I**) The human *MYC* transgene (*hg19MYC*) expression is largely restricted to the luminal compartment (**G**) and predominantly expressed in the VP (**H**), in accordance with the penetrance of prostatic intraepithelial neoplasia (**I**, PIN; mean ± SD; ^17^). WT: wild-type; VP: ventral prostate; DLP: dorsolateral prostate; AP: anterior prostate.

### Single-cell transcriptome delineates inter- and intra-prostate lobe heterogeneity

To determine key differences between murine prostate lobes, we projected the single-cell transcriptome data into the t-distributed stochastic neighbor embedding (tSNE) space. Using known markers (**Supplementary Figure S2A-B**), we identified nine major subpopulations of cells across prostate lobes (**Figure 1F**). Notably, basal cells (*Krt5*^+^, *Krt14*^Hi^) were the most abundant epithelial cell subtype observed in the AP and DLP lobes, whereas luminal cells (*Krt8*^Hi^, *Krt18*^Hi^) were overwhelmingly represented in the VP lobe. While murine *Myc* (*mm10Myc*) was expressed across all subpopulations and prostate lobes (**Supplementary Figure S2C and S3**), human *c-MYC* transgene expression (*hg19MYC*) was largely restricted to the luminal subpopulation (**Figure 1G**) and more prevalent in the VP lobe (**Figure 1H**), a feature in line with the greater penetrance of the MYC-driven PIN transformation observed in the VP lobe (**Figure 1I**)^17^.

The high representation of luminal cells coupled with a robust and uniform MYC-driven PIN transition in the VP enabled us to further define distinct luminal subpopulations. K-means clustering revealed a luminal subpopulation (*Krt8*^Hi^, *Krt18*^Hi^) common to both WT and MYC genotypes and characterized by high expression of *Krt4* but negative for *Nkx3-1* expression (*Krt4*^Hi^, *Nkx3-1*^-^; **Figure 2A-B** and **Supplementary Figure S4A**). Concurrent high expression of *Cd44, Tacstd2* (Trop2) and *Psca* suggests that this subpopulation corresponds to luminal progenitor cells ^18^. In untransformed VP, the main luminal cell cluster was composed of two subpopulations characterized by either high or low expression of androgen-responsive genes such as *Pbsn* and *Msmb* (**Supplementary Figure S4B**)^19,20^. Human *MYC* was predominately expressed in luminal cells (**Figure 2C-D**), resulting in an extensive transcriptional reprogramming within the luminal compartment (**Figure 2A-B**). Importantly, the distinct transcriptional profile of human *MYC* overexpressing luminal cells was identifiable even without inclusion of the human *MYC* transcript in the generation of the tSNE plot (**Supplementary Figure S5**). In agreement with MYC function in controlling transcriptional programs that favor cell growth and proliferation ^21^, we identified a subset of highly proliferative human *MYC* overexpressing luminal cells positive for cyclin B1, DNA topoisomerase II alpha and the marker of proliferation Ki-67 (*Ccnb1*^+^, *Top2a*^+^, *Mki67*^+^; **Figure 2B** and **Supplementary Figure S4C**), a state that was independent of human or murine *MYC* transcript levels (**Figure 2D)**. Finally, a limited number of cells belonging to hematopoietic (*Ptprc*^*+*^), vascular endothelium (*Pdgfra*^*+*^), smooth muscle (*Actg2*^*+*^) and adipocyte (*Fabp4*^*+*^) populations were also identified (**Figure 2B** and **Supplementary Figure S4D**). Taken together, these results demonstrate that MYC-driven transcriptional reprogramming can be readily captured by single-cell transcriptomics.

**Figure 2:**
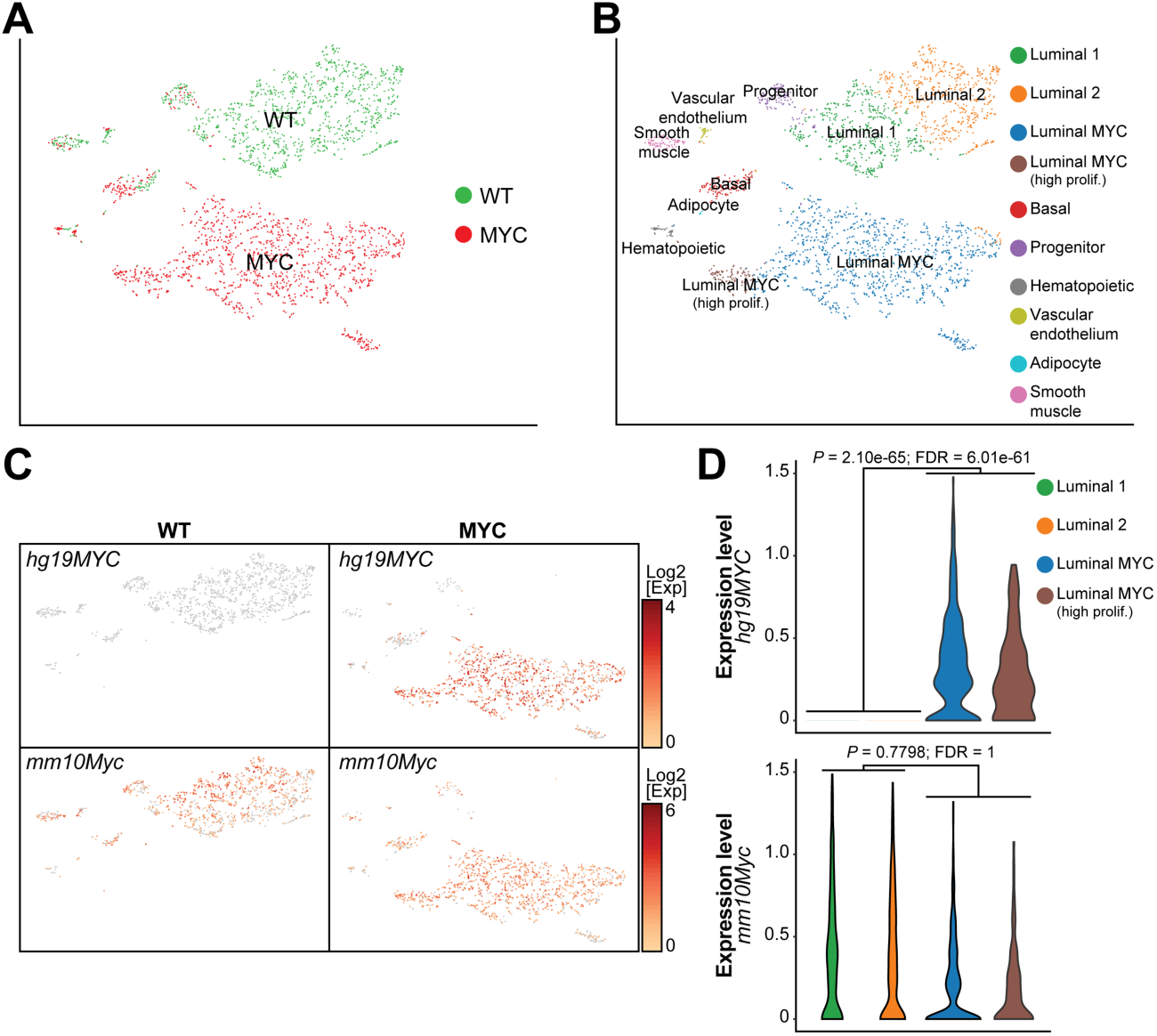
Single-cell transcriptome reveals distinct luminal cell subpopulations. (**A, B**) Single cell census of the WT and MYC-transformed VP (**A**) followed by unsupervised clustering revealed four luminal subsets (**B**). (**C, D**) Human *MYC* transcript (*hg19MYC*) is only observed in MYC-transformed VP and mostly restricted to the luminal subsets while murine *Myc* transcript (*mm10Myc*) is expressed across cellular populations and genotypes (**C**) and is not correlated with *hg19MYC* expression in luminal cells (**D**).

### MYC-driven luminal cells transformation dampens the AR transcriptional program

To define the transcriptional reprogramming driven by *MYC* overexpression in the VP lobe across cell subpopulations, we created a pseudobulk sample for each subpopulation and performed Gene Sets Enrichment Analyses (GSEA) using the Hallmark gene sets ^22^. As expected, the pseudobulk RNA-seq analysis showed that the MYC-driven transcriptional program enriched in gene sets related to cell proliferation (E2F_targets, G2M_chekpoint) or MYC-transcriptional activity *per se* (MYC_targets_V1/V2), was solely driven by the luminal cells (**Figure 3A-B**). In fact, the near totality of the MYC-driven transcriptional program captured by bulk RNA-seq is in line with the luminal cells transcriptional program. However, a large proportion of MYC-driven transcriptional reprogramming was undetected in bulk RNA-seq and only captured by single-cell transcriptomics. Notably, basal cells underwent an extensive transcriptional reprogramming (**Figure 3A**). Considering that human *MYC* transgene expression was detected in only a limited proportion of basal cells (18.3%; **Figure 2C**), this result suggests the existence of a paracrine transcriptional reprogramming upon MYC overexpression and prostate transformation. In addition, scRNA-seq revealed the downregulation of several transcriptional programs in luminal cells. Critically, the depletion of the Androgen_response gene set (**Figure 3A, C**), which was not accompanied with a global decreased in AR transcript and protein levels (**Figure 3D-E**; **Supplementary Figure S4E** and **Supplementary Data 1**), suggests a dampening of the AR transcriptional program driven by *MYC* overexpression as exemplified by loss of *Pbsn* and *Msmb* expression in the luminal compartment (**Supplementary Figure S4B**)^19,20^.

**Figure 3:**
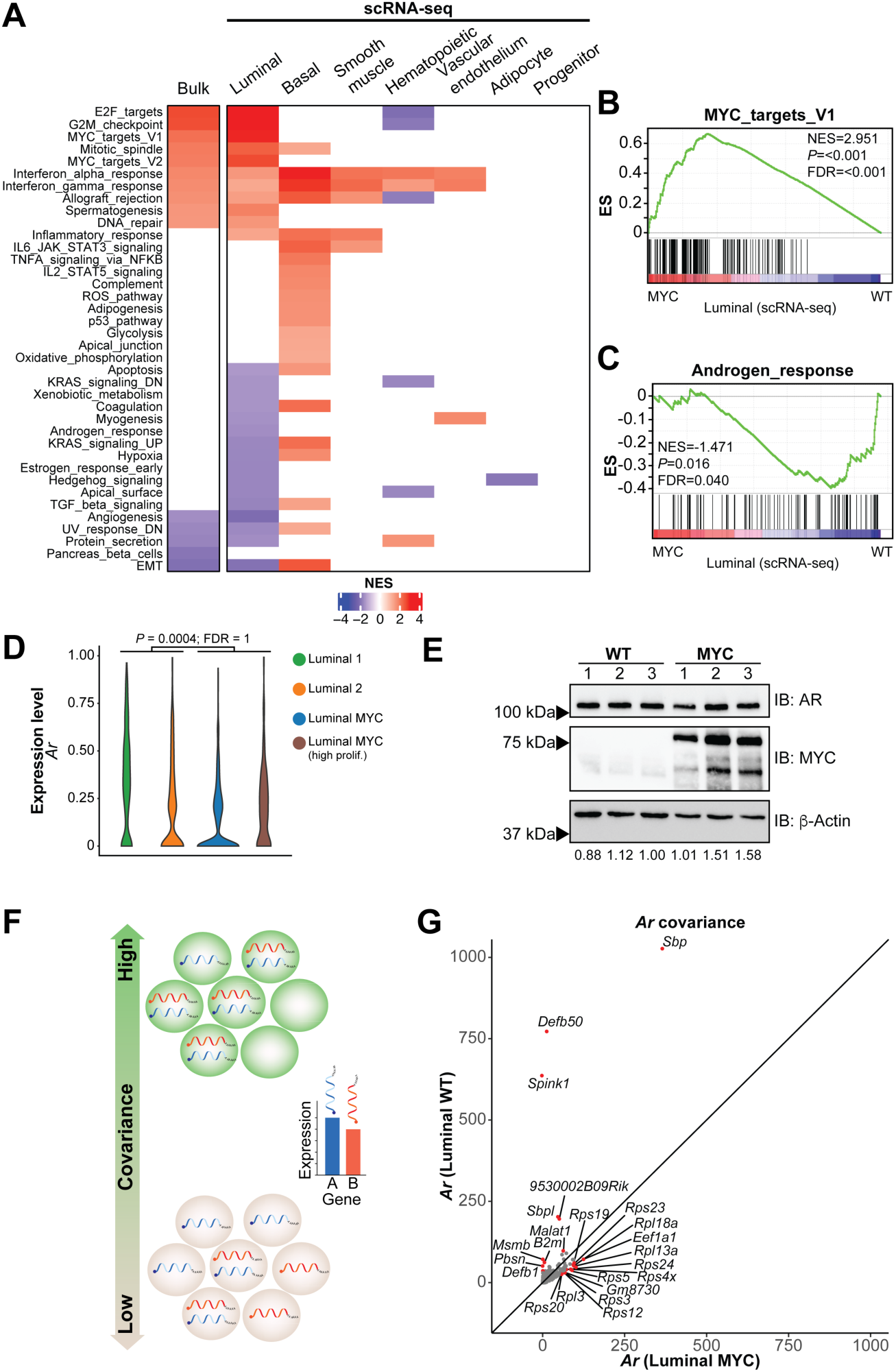
MYC-driven luminal cells transformation dampens the AR transcriptional program. (**A**) Gene Set Enrichment Analysis (GSEA, Hallmark, P<0.05 and FDR<0.1) revealed that the bulk RNA-seq transcriptional program associated with MYC overexpression is mostly driven by the luminal subset (matched scRNA-seq). (**B, C**) MYC overexpression is associated with an enriched MYC transcriptional program (**B**) and a depleted AR response (**C**) in the luminal subsets. (**D, E**) MYC overexpression does not alter AR transcript expression in the luminal compartment (**D**) and protein levels in the VP (**E**; numbers at the bottom represent AR levels relative to β-Actin). (**F**) Schematic representation of covariance analysis to determine co-expression (*i*.*e*. positive covariance) or mutually exclusive expression (*i*.*e*. negative covariance) between two genes at a single cell level. (**G**) Covariance analysis in the luminal subset reveals a shift from canonical AR target genes in the transcripts co-expressed with *Ar* upon MYC overexpression.

Thus, we sought to leverage single-cell transcriptomics to determine if *MYC* overexpression alters the nature of the transcripts co-expressed with *Ar* through a covariance analysis (**Figure 3F**)^23^. As expected, androgen-dependent genes such as *Pbsn, Msmb, Sbp, Defb50* and *B2m* or the prostate-specific *9530002B09Rik* were co-expressed with *Ar* in WT luminal cells (**Figure 3G**)^19,20,24-28^. Interestingly, both *Spink1* and *Malat1*, which are respectively associated with castration-resistant or enzalutamide-resistant disease ^29,30^, were strongly co-expressed with *Ar* only in untransformed tissues (**Figure 3G**), suggesting that these genes are also part of the normal androgen-dependent prostate epithelium homeostasis. Surprisingly, upon *MYC* overexpression, canonical AR target genes were no longer co-expressed with *Ar*. Instead, transcripts related to ribosome biogenesis, a key pathway driving cell growth and tumorigenesis and associated with MYC function ^31^, were co-expressed with *Ar* (**Figure 3G**). Altogether, these results indicate that AR-transcriptional program is compromised upon MYC overexpression.

### MYC overexpression alters the AR cistrome

To further characterize the mechanism whereby MYC overexpression negatively affects the AR-dependent transcriptional program, we utilized chromatin immunoprecipitation followed by high-throughput sequencing (ChIP-seq) to assess the AR cistrome. Although motif analysis of AR binding sites revealed the canonical androgen response element as the top enriched motif across genotypes (**Figure 4A**), unsupervised clustering uncovered a distinct AR cistrome driven by MYC overexpression (**Figure 4B**). Indeed, MYC overexpression resulted in a significant expansion of the AR cistrome with 1,695 sites gained compared to WT tissues (**Figure 4C**). Motif analyses revealed that AR gained sites are predominantly associated with the forkhead family of transcription factors motifs (forkhead response elements; FHRE), which includes the established regulator of AR transcriptional activity FOXA1, followed by androgen response elements (ARE; **Figure 4D**)^32^. Critically, FOXA1 occupancy was increased at AR gained binding sites in MYC-transformed prostate tissues compared to the WT counterpart (*P* = 2.23e-62; **Figure 4E** and **Supplementary Figure S6A**). Genomic regions gaining AR occupancy were characterized by increased histone H3K27 acetylation (H3K27ac; *P* = 4.39e-40; **Figure 4F**), a mark of active regulatory regions and transcriptional activity ^33^, supporting a differential usage of non-coding regulatory elements driven by AR in a MYC overexpressing context. To determine whether the repurposing of the AR cistrome upon MYC overexpression is associated with a distinct transcriptional program, we next integrated AR ChIP-seq to single-cell transcriptomics. Association of 1,695 AR binding sites gained upon MYC overexpression (**Figure 4C**) to the expression of nearby coding genes in the luminal cell subpopulations, ordered based on slingshot pseudotime inference across genotypes (**Supplementary Figure S6B**), highlighted three main expression patterns, namely a MYC-dependent increased, decreased or unchanged expression (**Figure 4G**). Using GSEA analysis and the Hallmark gene sets, we identified the MYC_targets_V1 as the top gene set enriched within the set of genes with increased expression. Conversely, we identified the Androgen_response among the gene sets that were significantly enriched within the set of genes with decreased expression (**Figure 4H**). Taken together these results indicate that, in the context of MYC overexpression, a reprogramming of the AR cistrome that drives an altered transcriptional program.

**Figure 4:**
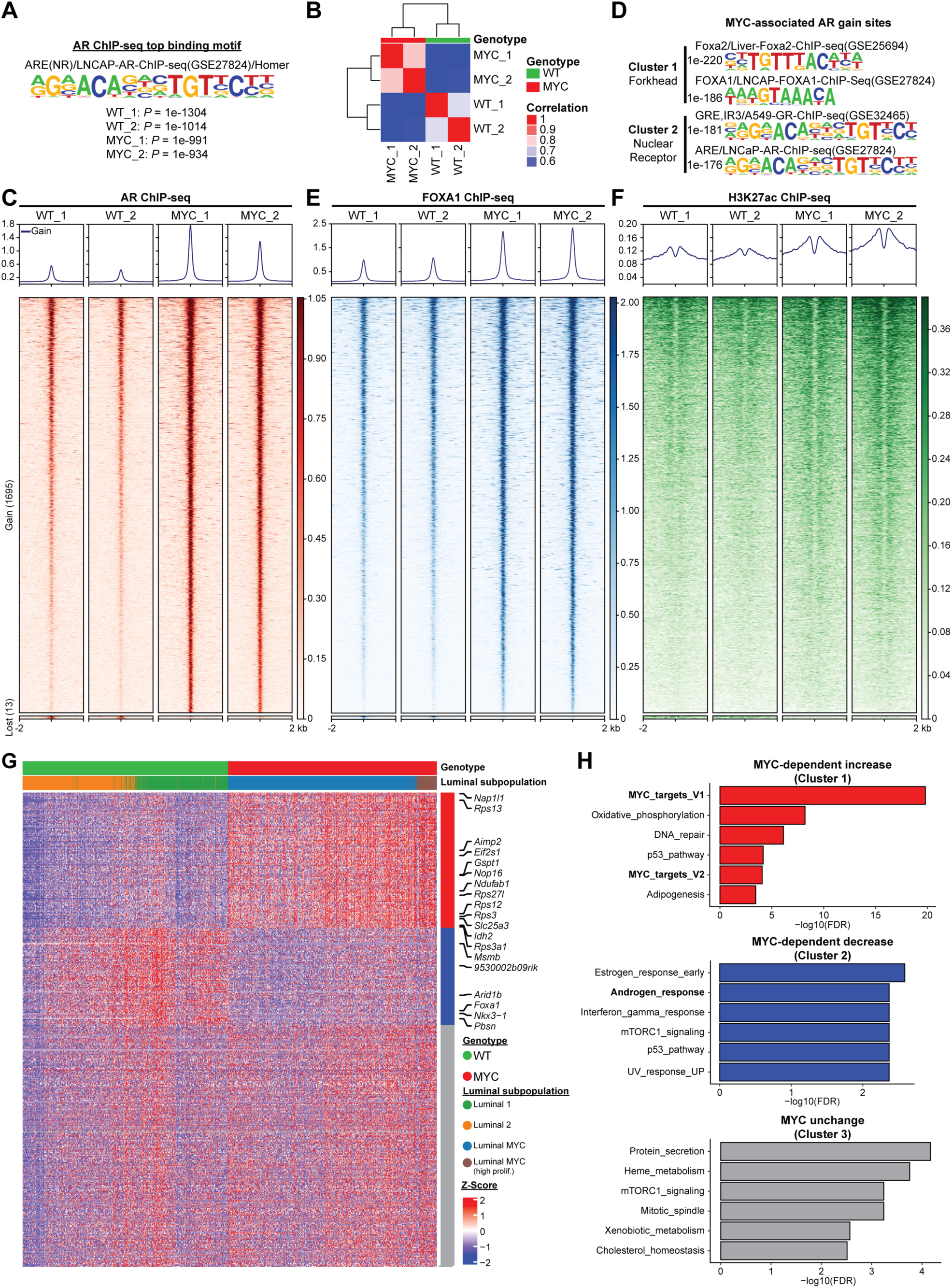
MYC overexpression alters the AR cistrome. (**A**) AR ChIP-seq identifies an androgen response element (ARE) as the top AR binding motif in WT and MYC-transformed VP. (**B**) Unsupervised pairwise correlation of the murine AR cistrome from all specimens. (**C**) MYC overexpression expands the AR cistrome as demonstrated by the heatmaps indicating AR binding intensity across 4 kb intervals. (**D**) Motif analysis of MYC-associated AR gained sites reveal forkhead response element (FHRE) and androgen response element (ARE). (**E, F**) AR gained sites are characterized by increased FOXA1 binding (**E**) and H3K27ac mark (**F**) in MYC-transformed VP. (**G**) Integration of the 1,695 AR bindings sites gained in MYC tumors with luminal single cell transcriptome grouped by k-means clustering (n=3). (**H**) GSEA analysis (Hallmark) revealed an enforced MYC transcriptional program (Cluster 1) and a diminished androgen response (Cluster 2) associated to MYC-dependent AR gained binding sites.

### Divergent MYC and AR transcriptional programs dictate disease progression

Since our results in the preclinical model uncovered a robust interplay between MYC and AR transcriptional programs, we next investigated whether this MYC-driven transcriptional reprogramming is clinically relevant. We used gene expression data to stratify 488 primary prostate cancer patients in the TCGA dataset based on the combined levels of the Hallmark Androgen_response (high; low) and MYC_targets_V1 (high; low) transcriptional signatures ^9^. Kaplan-Meier curves revealed that patients bearing a primary tumor characterized by divergent AR and MYC transcriptional programs experienced distinct rates of clinical progression. Tumors characterized by a low AR transcriptional signature with concurrent high MYC transcriptional signature (AR_low/MYC_high) were associated with the shortest time to biochemical recurrence (BCR) while tumors characterized by a high AR transcriptional signature with concurrent low MYC transcriptional signature (AR_high/MYC_low) were associated with the longest time to BCR (**Supplementary Figure S7A-B**). Interestingly, concordant AR and MYC transcriptional programs (AR_high/MYC_high; AR_low/MYC_low) were associated with an intermediate time to BCR (**Supplementary Figure S7A-B**). Recently, transcriptomic data from nearly 20,000 tumors revealed that patients bearing a localized treatment-naïve primary prostate cancer with low AR-activity (AR-A; based on a signature of nine canonical AR transcriptional targets) experience a shorter time to recurrence ^34^. Thus, we next sought to determine if MYC transcriptional activity status in low AR-A tumors could identify a more aggressive subtype of primary prostate cancer using the TCGA dataset. Strikingly, Kaplan-Meier curves revealed that it is the subset of low AR-A tumors with concurrent high MYC transcriptional signature that is associated with a faster time to BCR (AR_low/MYC_high *vs*. AR_low/MYC_low, *P* = 0.0001; **Figure 5A-B**). Importantly, we validated this finding in a previously published independent meta-analysis cohort combining 855 patients with individual patient-level data (**Figure 5C** and **Supplementary Figure S7C**)^35^.

**Figure 5:**
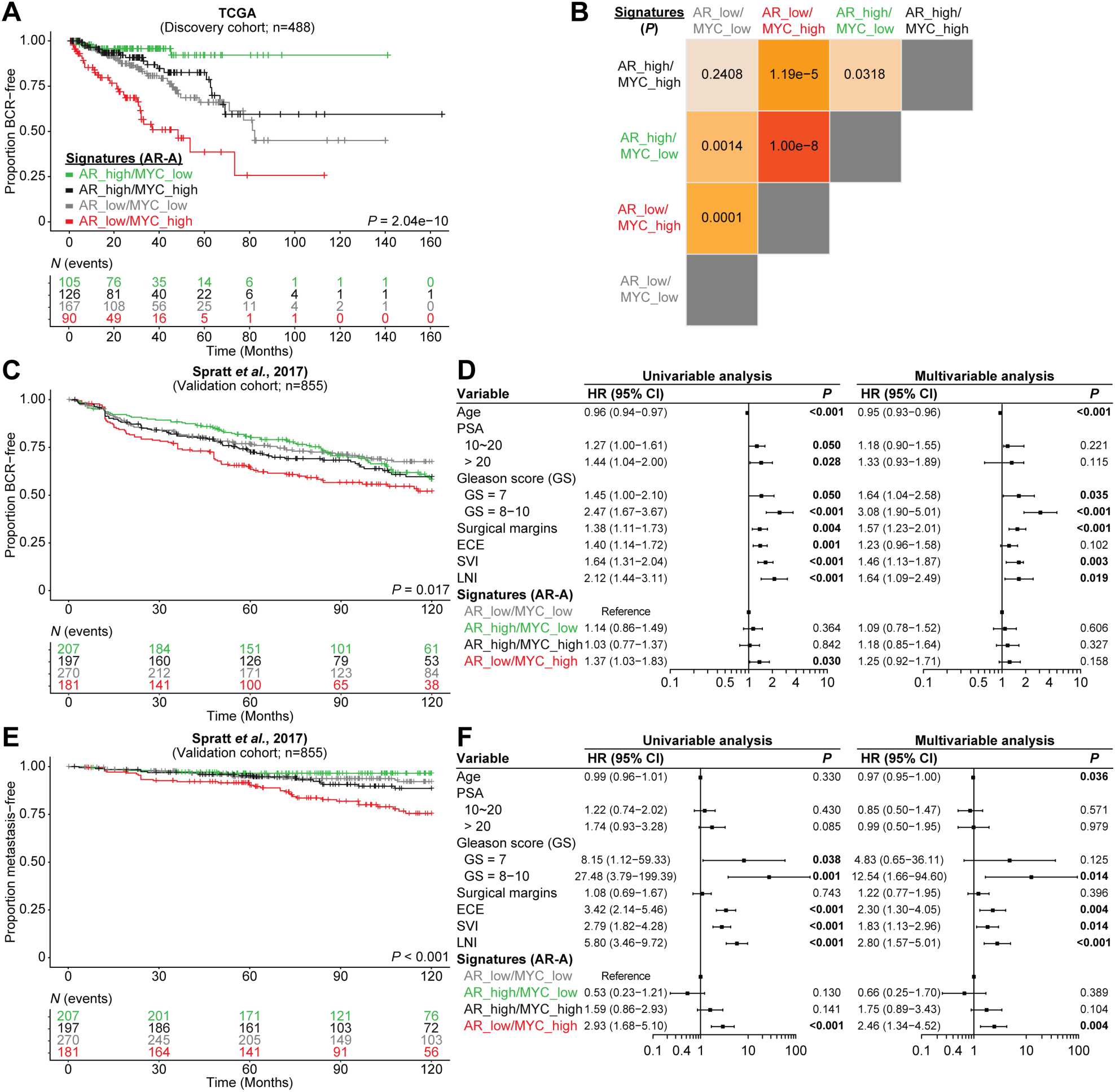
Divergent MYC and AR transcriptional programs dictate disease progression. (**A, B**) Kaplan-Meier curves (**A**) and log-rank tests (**B**) reveal that patients bearing a primary tumor characterized by low AR-activity (AR-A) and concurrent high MYC transcriptional signature (Hallmark) have a shorter time to biochemical recurrence (BCR) within the discovery cohort (TCGA). (**C, D**) Kaplan-Meier curves (**C**), univariable and multivariable analysis (**D**) confirms that tumors with concurrent low AR-A and high MYC transcriptional signatures develop BCR after radical prostatectomy more rapidly than low AR-A tumors without an active MYC transcriptional program in the validation cohort (Spratt *et al*., 2017). (**E, F**) Kaplan-Meier curves (**E**), univariable and multivariable analyses (**F**) reveal that tumors with concurrent low AR-A and high MYC transcriptional signatures are more likely to develop a metastatic disease. PSA: prostate-specific antigen; HR: hazard ratio; GS: Gleason score; ECE: extracapsular extension; SVI: seminal vesicles invasion; LNI: lymph node involvement.

Univariable analysis revealed that tumors with AR_low/MYC_high transcriptional signatures are associated with increased rates of BCR (Hazard Ratio (HR) = 1.37, 95% Confidence Interval (CI) 1.03-1.83; *P* = 0.030; **Figure 5D**), but this did not remain significant after adjusting for clinicopathologic risk factors in multivariable analysis (**Figure 5D** and **Supplementary Figure S7D**). Since low AR-A tumors were predicted to be less sensitive to androgen-deprivation therapy and more likely to develop metastatic disease after initial local therapy ^34^, we next asked whether a high MYC transcriptional activity allows for the identification of a more aggressive subtype of treatment-naïve primary prostate cancer. Strikingly, Kaplan-Meir curves revealed that patients with tumors harboring an AR_low/MYC_high signature were the most likely to develop metastatic disease (**Figure 5E** and **Supplementary Figure S7E**). Univariable analysis shows that AR_low/MYC_high tumors are associated with an increased risk to develop metastatic disease (HR = 2.93, 95% CI 1.68-5.10; *P* < 0.001; **Figure 5F** and **Supplementary Figure S7F**). Critically, this finding remained significant in a multivariable competing risks regression analysis adjusting for age, prostate-specific antigen (PSA), Gleason score, surgical margin status, extracapsular extension, seminal vesicles invasion and lymph node involvement (HR = 2.46, 95% CI 1.34-4.52; *P* = 0.004; **Figure 5F** and **Supplementary Figure S7F**). Altogether, our results suggest that concurrent AR_low/MYC_high transcriptional signatures identify a subgroup of patients that are predisposed to fail standard-of-care therapies and progress to develop metastatic disease.

### High *MYC* expression is associated with a dampened AR transcriptional program and an expanded AR cistrome in castration-resistant tumors

CRPC is characterized by MYC and AR amplification ^9,15,36^. Thus, we sought to assess the impact of MYC expression on the AR transcriptional program and cistrome. Gene expression profiling from 59 AR^+^ CRPC tumors revealed that AR activity is negatively correlated with *MYC* expression (**Figure 6A-B**)^37^. As expected, GSEA analysis revealed that *MYC*-high CRPC tumors are enriched for MYC transcriptional signatures. Strikingly, the Hallmark Androgen_response was the only gene set significantly depleted in *MYC*-high tumors (**Figure 6C**), supporting a role for MYC in dampening the canonical AR transcriptional program in the castration-resistant setting. We next determined whether this phenotype was associated with a repurposing of the AR cistrome using the LuCaP patient-derived xenografts (PDXs) series obtained from AR^+^ mCRPC samples (described in ^38^ and **Supplementary Figure S8A**). We selected eight specimens, for which the gene expression profiles were readily available, and stratified them into either the *MYC*-high or the *MYC*-low group based on transcript expression (**Figure 6D**)^37^. Importantly, AR transcript level was not different between the *MYC*-high and *MYC*-low groups (**Figure 6D**). Comparison of the AR cistrome between the two groups uncovered an alteration of AR binding in *MYC*-high mCRPC PDXs towards an expanded AR cistrome robustly associated with the forkhead family of transcription factors motifs (**Figure 6E-F, Supplementary Figure S8B**). Accordingly, greater FOXA1 occupancy was observed at AR gained binding sites in *MYC*-high compared to the *MYC*-low mCRPC PDXs (*P* = 1.74e-144; **Figure 6G** and **Supplementary Figure S8C**). These sites were also characterized by increased H3K27ac mark (*P* = 3.54e-268; **Figure 6H**), in agreement with the MYC-driven murine prostate cancer model (**Figure 4**). Critically, differential AR chromatin occupancy between both groups was associated with a dampened AR transcriptional program in the *MYC*-high group (**Figure 6I**). Taken together, these results support the existence of a distinct AR cistrome in *MYC* overexpressing CRPC associated with a diminished AR transcriptional program.

**Figure 6:**
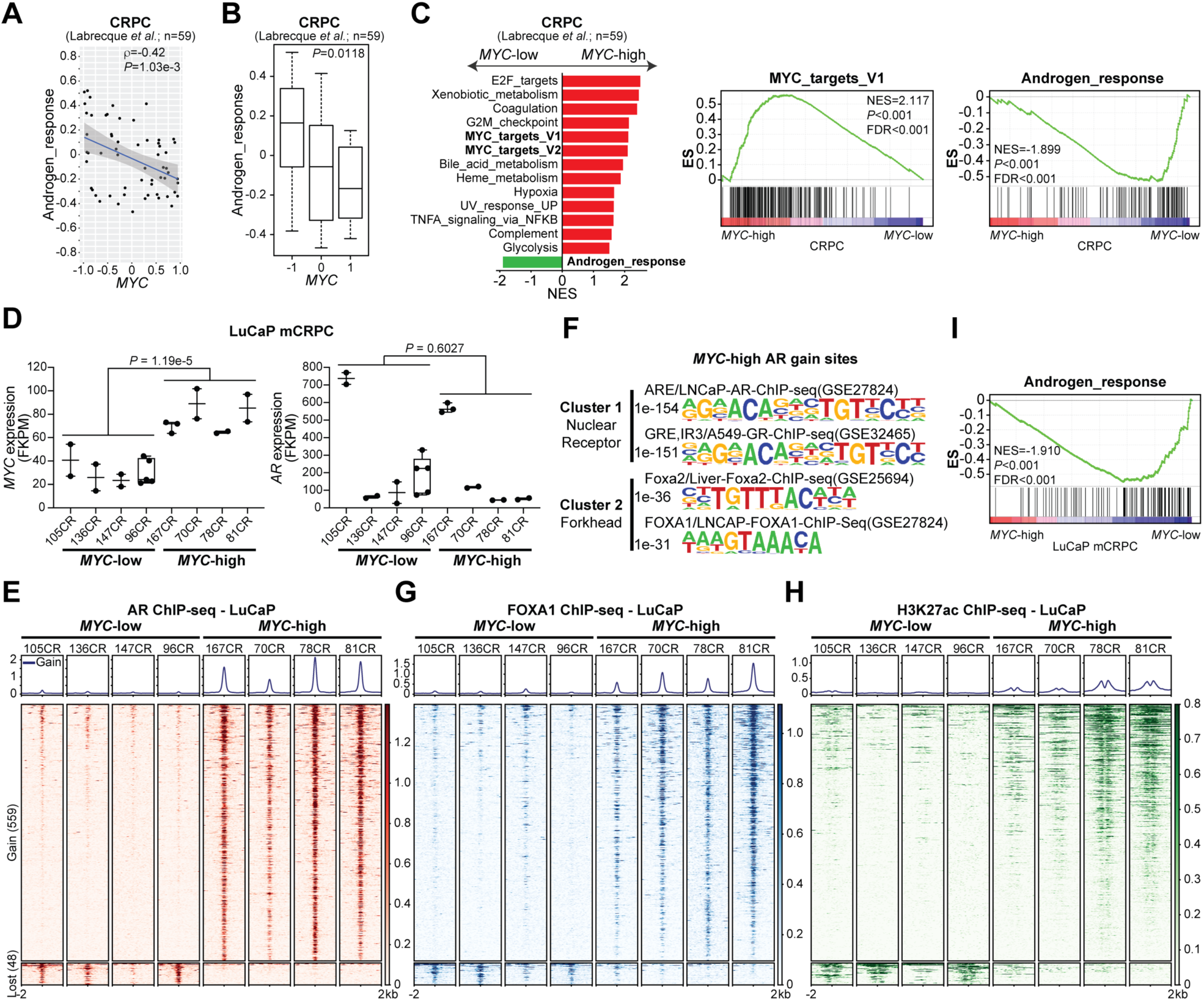
High *MYC* expression is associated with a dampened AR transcriptional program and an expanded AR cistrome in castration-resistant tumors. (**A, B**) AR activity is inversely correlated with *MYC* expression in CRPC clinical samples (**A**) and significantly lower in *MYC*-high tumors (**B**; median ± 1.5 interquartile range). (**C**) Gene Set Enrichment Analysis (GSEA, Hallmark, P<0.05 and FDR<0.1) revealed an enriched MYC transcriptional program and a depleted AR response in *MYC*-high CRPC. (**D, E**) *MYC*-high mCRPC LuCaP patient-derived xenografts (PDXs) have similar levels of *AR* (**D**; median ± min to max) but are associated with an expanded AR cistrome as demonstrated by the increased binding intensity across 4 kb intervals at AR gained sites (**E**). (**F**) Motif analysis of *MYC*-associated AR gained sites reveal ARE and FHRE. (**G, H**) AR gained sites are characterized by increased FOXA1 binding (**G**) and H3K27ac mark (**H**) in *MYC*-high mCRPC LuCaP. (**I**) AR cistrome in *MYC*-high mCRPC LuCaP PDXs is associated with a diminished androgen response.

### MYC overexpression disrupts the AR transcriptional program by pausing AR regulated genes

To assess for direct effects of AR in mediating this transcriptional reprogramming we leveraged the preclinical model of MYC-driven prostate cancer and performed binding and expression target analysis (BETA) to integrate MYC-driven gene expression changes in murine VP with genome-wide AR binding data ^39^. This analysis revealed that AR binding was significantly associated with genes downregulated by MYC overexpression (*P* = 2.32e-5; **Figure 7A**). Along this line, AR binding was found to be increased at genomic regions nearby Androgen_response genes alongside the H3K27ac mark following MYC overexpression (**Figure 7B-C**), in contrast with the accompanied depletion of the Androgen_response gene set (**Figure 3C**). For example, AR and FOXA1 binding was increased in the promoter region of *Pbsn* (**Figure 7D**), an AR-dependent gene whose transcript and protein levels were both severely downregulated following MYC overexpression (**Figure 7E-F**; **Supplementary Figure S4B** and **Supplementary Data 2**). In the promoter region of *Msmb*, another AR-dependent gene previously characterized as a tumor suppressor ^40^, AR and FOXA1 binding as well as the H3K27ac mark levels were maintained although *Msmb* transcript levels were also downregulated by MYC overexpression (**Figure 7G-H** and **Supplementary Figure S4B**). These results suggest that MYC-driven repression of the AR transcriptional program is not associated with a disengagement of AR or the loss of the H3K27ac mark.

**Figure 7:**
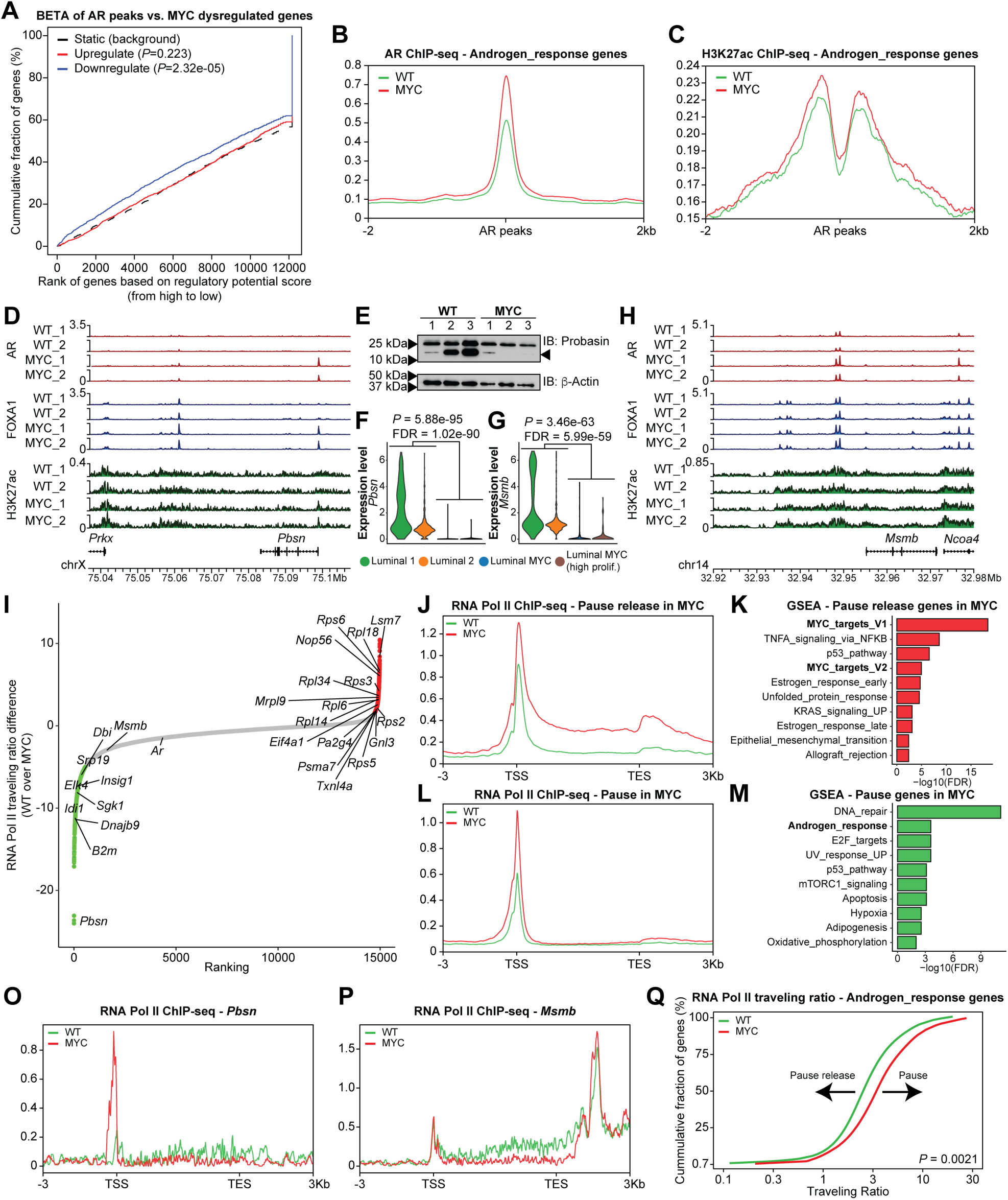
MYC overexpression disrupts the AR transcriptional program by pausing AR regulated genes. (**A**) BETA analysis revealed that AR binding sites are associated with gene downregulation following MYC overexpression. (**B, C**) Despite a dampened AR transcriptional program, higher levels of the AR binding (**B**) and H3K27ac mark (**C**) are observed nearby AR response genes. (**D-F**) AR, FOXA1 and H3K27ac tracks at *Pbsn* locus, an AR-dependent gene, reveal unchanged or heightened AR and FOXA1 binding (**D**) albeit decreased protein (**E**) and transcript levels (**F**) following MYC overexpression. (**G, H**) Unchanged AR and FOXA1 binding and H3K27ac mark at *Mmsb* locus (**G**), an AR-dependent gene downregulated by MYC overexpression (**H**). (**I**) RNA Pol II traveling ratio differences following MYC overexpression in murine VP. (**J, K**) Pause release genes following MYC overexpression are characterized by greater RNA Pol II occupancy at gene body (**J**) and are enriched for MYC transcriptional signatures (GSEA, Hallmark, P<0.05 and FDR<0.1; **K**). (**L, M**) Pause genes following MYC overexpression are characterized by greater promoter-proximal RNA Pol II occupancy (**L**) and are enriched for AR transcriptional signature (GSEA, Hallmark, P<0.05 and FDR<0.1; **M**). (**O, P**) Increased RNA Pol II occupancy at the promoter of *Pbsn* (**O**) and decreased occupancy at the gene body of *Msmb* (**P**) following MYC overexpression. (**Q**) RNA Pol II traveling ratio reveals greater promoter-proximal pausing at Androgen_response genes.

Using the androgen responsive LNCaP prostate cancer cell line, Barfeld and colleagues have previously reported that MYC overexpression antagonizes the transcriptional activity of the AR ^15^. Similarly to the MYC-driven genetically engineered prostate cancer mouse model, MYC overexpression in LNCaP cells was associated with the depletion of the Hallmark Androgen_response gene set (**Supplementary Figure S9A**). Annotation of the AR cistrome and gene expression data by BETA revealed that AR binding is associated with downregulated genes, supporting a global reduction in AR transcriptional activity driven by MYC overexpression. Conversely, MYC cistrome was predominantly associated with upregulated genes, consistent with its role as a transcriptional activator (**Supplementary Figure S9B**). Again, AR binding nearby Androgen_response genes remained largely unchanged following MYC overexpression. Interestingly, MYC binding nearby MYC_targets_V1 genes also remained unchanged following MYC overexpression despite a significant enrichment of the MYC_targets_V1 gene set (**Supplementary Figure S9C**). Inspection of AR and MYC binding in the vicinity of canonical AR-dependent genes such as *KLK3* and *TMPRSS2* also revealed unchanged binding profiles (**Supplementary Figure S9D**).

Based on the evidence for MYC regulation of RNA Pol II pause release ^41^, we leveraged RNA Pol II ChIP-seq to determine genome-wide RNA Pol II traveling ratio (*i*.*e*. RNA Pol II density in the promoter-proximal region over the RNA Pol II density in the transcribed region) *in vivo* following MYC overexpression in murine VP (**Figure 7I**). As expected, genes with reduced RNA Pol II traveling ratio following MYC overexpression were enriched for MYC transcriptional signatures, indicative of pause release at these sites (**Figure 7J-K** and **Supplementary Figure S10A**). Critically, genes with greater RNA Pol II traveling ratio were enriched for the AR transcriptional signature, suggestive of enhanced RNA Pol II pausing at AR-regulated genes (**Figure 7L-M** and **Supplementary Figure S10A**). Along this line, ChIP-seq revealed a build-up of RNA Pol II occupancy at the promoter of the AR-regulated gene *Pbsn* following MYC overexpression (**Figure 7O**). At the *Msmb* locus, another AR-regulated gene, RNA Pol II occupancy remained unchanged at the promoter region but was abrogated at the gene body in the MYC overexpressing condition (**Figure 7P**). These features are in stark contrast to MYC-regulated genes such as *Rps3* and *Rps5* for which we observed an increase RNA Pol II occupancy at the gene body in the MYC overexpressing condition (**Supplementary Figure S10B-C**). Since these patterns suggest a MYC-driven altered ratio of initiating and elongating RNA Pol II at AR-regulated genes, we next determined the RNA Pol II traveling ratio at Androgen_response genes. Strikingly, RNA Pol II traveling ratio at Androgen_response genes was significantly increased by MYC overexpression (*P* = 0.0021; **Figure 7Q** and **Supplementary Figure S10D**), supporting MYC-driven RNA Pol II promoter-proximal pausing and consequently non-productive transcription at AR-dependent genes. Altogether these findings support RNA Pol II promoter-proximal pausing as a potential mechanism for MYC-mediated transcriptional repression at AR regulated genes associated with the canonical AR transcriptional signature (**Graphical Summary**)^42^.

## DISCUSSION

In this study, we report the impact of MYC overexpression *in vivo* on the AR transcriptional program. By leveraging the expression of a human MYC transgene (*hg19MYC*) observed at a single-cell level in murine prostatic tissues, our data demonstrate that MYC overexpression robustly reprograms luminal (*Krt8*^Hi^, *Krt18*^Hi^) cells toward a repressed AR transcriptional program, a feature contrasting with the supporting role of MYC on the AR transcriptional program in the apocrine breast cancer subtype ^43^. Our single-cell transcriptome data delineate a minor luminal subpopulation expressing high levels of *Cd44, Tacstd2* (Trop2) and *Psca* markers associated with luminal progenitor cells ^18^. Recently, single-cell transcriptomics performed in the murine AP lobe also revealed a distinct but rare luminal subpopulation anatomically lining the proximal duct and expressing *Tacstd2* (Trop2), *Psca* as well as *Ly6a* (Sca-1), *Krt4* and *Cldn10* ^44^. An independent study suggested that the luminal subpopulation expressing high levels of progenitor markers such as *Tacstd2* (Trop2), *Psca, Ly6a* (Sca-1) and *Krt4* corresponds to urethral luminal cells extending into the proximal ducts of the prostate ^45^. Since the luminal progenitor population identified in the VP lobe expressed all the aforementioned markers (**Supplementary Figure S4A, F**), we cannot rule out the possibility that they might be of urethral origin. Regardless, these progenitor cells were not transcriptionally reprogramed following MYC overexpression (**Figure 3A**).

In analyzing the expression of *hg19MYC* transcript driven by the ARR_2_Pb promoter we found it was not detected in WT prostates, as expected. Surprisingly, we detected low, but consistent *hg19MYC* expression in non-luminal subpopulations (basal: 17/93 (18.3%); hematopoietic: 3/35 (8.6%); vascular endothelium: 1/8 (12.5%); **Figure 2C**). While the ARR_2_Pb promoter used to drive *hg19MYC* expression has been described as highly specific for prostatic epithelium ^14,20,46^, our single-cell transcriptome highlights a potentially underappreciated leaky expression of ARR_2_Pb-driven transgene. However, these seemingly stochastic events are likely transient since Hi-MYC mice do not develop other MYC-driven malignancies, such as B-cell leukemia/lymphoma ^47^. With the increasing availability of single-cell transcriptomic profiles from various genetically engineered mouse models (GEMMs), it is expected that tissue specific promoter specificity will be reassessed through a new lens.

MYC is commonly amplified in primary prostate cancer and is overexpressed in 37% of metastatic disease ^9,48^. Considering that prostate cancer cells that develop resistance to AR-targeted therapy usually maintain AR expression ^49,50^, the interplay between MYC and AR is likely to remain critical as the disease progress to the CRPC stage. Importantly, our analyses exposed a subtype of primary prostate cancer characterized by divergent AR (low) and MYC (high) transcriptional signatures that are predisposed to fail standard-of-care therapies and progress to the mCRPC stage (**Figure 5**). Arriaga and colleagues have recently reported a MYC and RAS co-activation signature associated with metastatic progression and failure to anti-androgen treatments ^51^. It is thus tempting to speculate that MYC decreases the reliance of prostate cancer cells on the canonical AR transcriptional program, therefore facilitating resistance to AR-targeted therapies. Along this line, Bai *et al*. recently showed that a c-Myc inhibitor disrupting c-Myc and Max dimerization sensitizes enzalutamide-resistant prostate cancer cells to growth inhibition by enzalutamide ^52^. Considering that transition from CRPC to neuroendocrine prostate cancer (NEPC) is driven by N-Myc, which also abrogates AR transcriptional program, and that N-Myc is functionally complementary to c-Myc in various processes ^53,54^, it is now evident that Myc family members are key to prostate cancer etiology and resistance to standard-of-care therapies.

Intriguingly, although MYC overexpression antagonizes the AR transcriptional program, this was not associated with a diminished but rather an expanded AR cistrome, characterized by FOXA1 co-occupancy and an active chromatin state. Data from our MYC-driven prostate cancer mouse model, together with a previously published LNCaP model engineered to overexpress MYC, revealed that MYC-driven repression of the AR transcriptional program is not associated with a disengagement of AR or the loss of the H3K27ac mark. Rather, we observed greater RNA Pol II promoter-proximal pausing and non-productive transcription at AR-dependent genes repressed by MYC *in vivo*. Importantly, no evidence of direct interaction between MYC and AR has been found ^15,52^, suggesting that the suppression of the AR transcriptional program is not guided by a physical interaction with MYC but rather by a MYC-induced RNA Pol II pausing overcoming the AR enhancers driving AR-regulated genes. Taken together, these results support cofactor redistribution driven by increased MYC expression and resulting in greater RNA Pol II promoter-proximal pausing as a potential mechanism for MYC-mediated transcriptional repression at genes regulated specifically by the AR (**Graphical Summary**)^42,55^.

Altogether, our study revealed an intricate crosstalk between the AR, MYC, FOXA1 and RNA Pol II resulting in a corrupted AR transcriptional program and promoting prostate cancer initiation and progression to the mCRPC stage. Considering that a simple dietary intervention meant to reduce saturated fat consumption can dampen MYC transcriptional program, and the recent development of viable MYC inhibitors for therapeutic interventions ^17,56^, we foresee that targeting MYC may help restore a canonical AR transcriptional program and sensitize prostate cancer to AR-targeted therapies.

## Acknowledgements

We thank Zach Herbert for technical assistance, Noriko Uetani for graphical summary design and drawings and Marie-Claude Gingras and Livia Garzia for critical review of this manuscript. J.L. is a recipient of a Canadian Institute of Health Research Frederick Banting and Charles Best Canada Graduate Scholarship-Master’s and of a Research Institute of the McGill University Health Centre M.Sc. Studentship award. Establishment and characterization of the LuCaP PDX models has been supported by the Pacific Northwest Prostate Cancer SPORE (P50CA97186), the U.S. Department of Defense Prostate Cancer Biorepository Network (W81XWH-14-2-0183), the National Cancer Institute (P01 CA163227), the Prostate Cancer Foundation, the Institute for Prostate Cancer Research, and the Richard M. Lucas Foundation. We would like to thank the patients who generously donated tissue that made this research possible. G.Z. is a recipient of an Idea Development Award from the U.S. Department of Defense (PC150263) and the Barr Award from the Dana-Farber Cancer Institute. This work has been supported by National Institutes of Health grants to K.W.W. (R01 CA238039; R01 CA251599), a Prostate Cancer Foundation Challenge Award to M.M.P. and M.L.F. and grants to M.L.F. (National Institutes of Health, R01 GM107427 and R01 CA193910; U.S. Department of Defense grant, W81XWH-19-1-0565; the H.L. Snyder Medical Research Foundation). E.C., M.B. and H.W.L. acknowledge support from the National Institutes of Health (P01 CA163227-06A1). D.P.L is a Lewis Katz – Young Investigator of the Prostate Cancer Foundation and is the recipient of a Scholarship for the Next Generation of Scientists from the Cancer Research Society and is also a Research Scholar – Junior 1 from The Fonds de Recherche du Québec – Santé. The work reported here was funded by a Canadian Institutes of Health Research project grant (PJT-162246) to D.P.L.

## Author contributions

Conceptualization, X.Q., M.B., H.W.L. and D.P.L.;

Methodology: X.Q., N.B., A.F. and Y.X.;

Software: X.Q., N.B., A.F. and Y.X.;

Validation: Y.L., E.D. and D.E.S.;

Formal Analysis, X.Q., N.B., A.F., Y.X., S.G., Q.T., Y.Z. and D.P.L.;

Investigation, T.H., A.d.P., A.M.L., J.L., G.Z., S.S., J-H.S., C.B., E.O’C., P.C. and D.P.L.;

Resources, E.M.S., R.J.K., S.W., L.E., M.L., K.W.W., M.M.P., E.C., M.L.F., X.S.L., M.B., H.W.L. and D.P.L.;

Data Curation, X.Q. and N.B.;

Writing – Original Draft, D.P.L.;

Writing – Review & Editing, X.Q., N.B., G.Z., E.C., M.F., H.W.L. and D.P.L.;

Visualization, X.Q., N.B. and D.P.L.;

Supervision, H.W.L. and D.P.L.;

Project Administration, H.W.L. and D.P.L.;

Funding Acquisition, M.B., H.W.L. and D.P.L.

## METHODS

### Animal husbandry

FVB Hi-MYC mice (strain number 01XK8), expressing the human c-MYC transgene in prostatic epithelium, were obtained from the National Cancer Institute Mouse Repository at Frederick National Laboratory for Cancer Research ^14^. Upon weaning (3 weeks), male mice heterozygous for the transgene (MYC), together with their wild type littermates (WT), were fed a purified diet (TD.130838, Envigo). Animals were kept on a 12-hour light / 12-hour dark cycle, and allowed free access to food and water at the Dana-Farber Cancer Institute (DFCI) Animal Resources Facility. The animal protocol was reviewed and approved by the DFCI Institutional Care and Use Committee (IACUC), and was in accordance with the Animal Welfare Act. For protein expression experiments, mice were housed in the Animal Resources Facility at the Research Institute of the McGill University Health Centre (RI-MUHC) where they were fed a regular lab chow (T.2918, Envigo) from the time of weaning. The animal protocol followed the ethical guidelines of the Canadian Council on Animal Care, and was approved by the RI-MUHC Glen Facility Animal Care Committee (FACC).

#### Genotyping

Tail snips were sent to Transnetyx (Transnetyx, Inc.) for genotyping or genomic DNA was extracted from ear punches using 0.4 mL of lysis buffer (100 mM Tris-HCl pH 7.5, EDTA 5 mM, 2% SDS, 200 mM NaCl and 100 µg/µL freshly added Proteinase K). Samples were incubated overnight at 52°C. After centrifugation at 13,000 rpm for 20 minutes, the supernatant was collected and mixed by inversion with 0.4 mL isopropanol to precipitate the DNA, which was pelleted by centrifugation for 5 minutes, then washed with 0.5 mL 70% ethanol and dissolved in 10 µL molecular grade water. The presence of the MYC transgene was detected by polymerase chain reaction (PCR), using the following primer combination: primer 1: 5’ AAA CAT GAT GAC TAC CAA GCT TGG C 3’and primer 2: 5’ ATG ATA GCA TCT TGT TCT TAG TCT TTT TCT TAA TAG GG 3’. PCR products were resolved using a 2% agarose tris-acetate-EDTA gel and a 177 bp band was visualized using the ChemiDoc™ imaging system (Bio-Rad).

### Tissue specimens

#### FVB Hi-MYC model

At 12 weeks of age, mice were euthanized by CO_2_ / isoflurane followed by cervical dislocation. Mouse prostate lobes (AP, DLP, VP) were dissected, weighed and immediately processed for bulk and single-cell transcriptomics or flash-frozen in liquid nitrogen for chromatin immunoprecipitation or protein expression experiments. Tissues were consistently collected during the same periods to minimize inter-samples and circadian rhythm variability.

#### mCRPC LuCaP PDXs

Informed consent was obtained to collect human mCRPC tissues and generate the patient-derived xenograft tumors as described previously ^37,38^. The study was approved by the University of Washington Human Subjects Division institutional review board (no. 39053). All animal studies were approved by University of Washington IACUC and performed according to NIH guidelines. Molecular characterization of AR^+^ mCRPC LuCaP PDXs 70CR, 78CR, 81CR, 96CR, 105CR, 136CR and 147CR was previously described ^37,38^. LuCaP PDX 167CR was established from a liver metastasis of 77-year-old Caucasian male who died of abiraterone-, carboplatin- and docetaxel-resistant CRPC. LuCaP 167CR expresses AR, responds to castration and is negative for synaptophysin. PDX cellular morphology recapitulates the original liver metastasis (**Supplementary Figure S8A**).

### Bulk RNA-sequencing

#### FVB Hi-MYC model

Fresh prostate lobes from 12-week-old mice were dissociated to form a single cell suspension. Prostate lobes were minced with a sterile razor blade and resuspended in collagenase/hyaluronidase (#07912, Stemcell Technologies) diluted in DMEM/F-12 (#36254, Stemcell Technologies) at 37°C for 2 hours. After dissociation, cells were centrifuged (350 x g for 5 minutes) and resuspended in 5 mL of prewarmed 0.25% trypsin/EDTA (#07901, Stemcell Technologies) at 37°C for 5 minutes. Trypsinization was stopped with 10 mL of cold HBSS (#37150, Stemcell) supplemented with 2% of regular cell culture grade FBS. Cells were centrifuged (350 x g for 5 minutes) and resuspended in 1 mL of prewarmed dispase (#07913, Stemcell Technologies) and 100 μL of DNase I (#07900, Stemcell Technologies) and passed 5 times through a 27G syringe needle. Cells were then mixed with 10 mL of cold HBSS supplemented with 2% FBS, filtered through a 40 μm cell strainer (#27305, Stemcell Technologies), centrifuged (350 x g for 10 minutes) and resuspended in PBS. RNA from an equal number of cells was extracted using the miRNeasy Micro Kit (#217084, Qiagen) coupled with on-column DNAse treatment (#79254, Qiagen). RNA sample concentration was measured and subjected to quality evaluation, using a Bioanalyzer RNA 6000 Nano kit (#5067-1511, Agilent). The Dana-Farber Cancer Institute Molecular Biology Core Facilities prepared libraries from 500 ng of purified total RNA, using TruSeq Stranded mRNA sample preparation kits (#RS-122-2101, Illumina) according to the manufacturer’s protocol. Finished libraries were quantified by the Qubit dsDNA High-Sensitivity Assay Kit (#32854, Thermo Fisher Scientific), by an Agilent TapeStation 2200 system using D1000 ScreenTape (#5067-5582, Agilent), and by RT-qPCR using the KAPA library quantification kit (#KK4835, Kapa Biosystems), according to the manufacturers’ protocols; pooled uniquely indexed RNA-seq libraries in equimolar ratios were sequenced to a target depth of 40M reads on an Illumina NextSeq500 run with single-end 75 bp reads. Read alignment, quality control and data analysis was performed using VIPER ^57^, RNA-seq reads were mapped by STAR ^58^ and read counts for each gene were generated by Cufflinks ^59^. Differential gene expression analyses were performed on absolute gene counts for RNA-seq data and raw read counts for transcriptomic profiling data using DESeq2 ^60^.

#### mCRPC LuCaP PDXs

LuCaP PDX tumor samples were collected from castrated CB 17 SCID male mice. Frozen tumors were used for RNA extraction and RNA-seq analysis as described previously ^37^.

#### LNCaP MYC model

Published gene expression data (GSE73995; ^15^) was downloaded and reanalyzed.

### Single-cell RNA-sequencing

Cell preparation for 3’ barcoded scRNA-seq (#120237, Chromium V2 assay) was performed according to the manufacturer’s protocol (10X Genomics) targeting 5000 cells from single-cell suspensions of freshly processed prostate lobes as described above. Single-cell RNA-seq data were preprocessed using the 10x genomics Cell Ranger (https://www.10xgenomics.com/) to obtain the UMI (unique molecular identifier) counts for each gene. To get a reliable single cell transcriptome dataset, we excluded the cells with fewer than 200 genes expressed (UMI > 0) or the cells with more than 80% UMIs from mitochondrial genes. The filtered data was then normalized and scaled by using Seurat to remove unwanted sources of variations ^61^. tSNE was performed on the normalized data to visualize the single cells in two-dimensional space by using the result of principal component analysis (PCA). Unsupervised clustering was performed by using the “FindClusters” function in the Seurat package with parameters of resolution = 0.8. Genes with differential expression between clusters were obtained by using Wilcoxon rank-sum test. FDR was calculated to correct for multiple testing.

#### Specific gene expression levels

The normalized expression level for all cells was calculated by the Seurat R package (3.1.1). The Violin plots were created by the geom_violin function in ggplot2 (3.3.2), scale option set to ‘area’.

#### Covariance analysis

The covariance for all genes with *Ar* is calculated by the cov function in stats R package (3.6.0). Genes that have covariance difference larger than 30 between the WT and MYC samples were colored in red and labeled in the plot.

#### Slingshot pseudotime inference

Pseudotime inference is done by the slingshot version 1.3.1. K-means clustering results and tSNE coordinates were used as input for the pseudotime inference.

### Bioinformatics analyses – bulk RNA-seq and scRNA-seq

#### Bulk RNA-seq and scRNA-seq gene expression correlation

X-axis is the log(scRNA-seq sum of UMI from all cells), Y-axis is log(bulk RNA–seq – raw read counts). Correlation is calculated based on Pearson Correlation. The Venn diagram is the overlap expressed genes between scRNA-seq and bulk RNA-seq. A gene is considered as expressed when the sum of UMI from all cells is larger than 0 in scRNA-seq or raw read counts is larger than 0 in bulk-RNA-seq.

#### Sample-sample correlation and principal component analysis (PCA)

Sum of UMI from all cells in scRNA-seq and raw read counts in bulk RNA-seq for matched samples were calculated. Batch effects between scRNA-seq and bulk RNA-seq data were removed using the ComBat approach from SVA (3.18.00). Pearson correlation and principal components were calculated using the counts after removal of batch effect.

#### Gene set enrichment analysis (GSEA)

All GSEA were done using pre-ranked analysis (GSEA Java; v4.1.0) with Hallmark gene sets (<u>h.all.v7.2.symbols.gmt</u>). Heatmap visualization of normalized enrichment score (NES) was obtained using ComplexHeatmap (2.2.0) R package ^62^.

### Protein expression

Fresh-frozen VP tissues from 12-week-old mice were sliced on ice with stainless steel disposable scalpels (Fisher Scientific) then homogenized in RIPA buffer (20 mM Tris-HCl pH 7.5, 150 mM NaCl, 1 mM EDTA, 1% TRITON-X) supplemented with phosphatases and protease inhibitors (Mini, Pierce™, Thermo Fisher) using a tissue grinder kit (Kontes). Equal amounts of protein (15 µg; Pierce™ Rapid Gold BCA Protein Assay, Thermo Fisher) were resolved on 8-12% Tris-glycine SDS-polyacrylamide gels and transferred to nitrocellulose blotting membranes (Bio-Rad), following standard procedures. Membranes were probed with the following antibodies according to the manufacturer’s instructions: rabbit monoclonal [Y69] anti-c-MYC (#ab32072, Abcam), rabbit monoclonal [ER179(2)] anti-AR (#ab108341, Abcam), mouse monoclonal [F6] anti-probasin (#sc-393830, Santa Cruz Biotechnology) or rabbit polyclonal anti-β-Actin (#4967, Cell Signaling Technology). Densitometry analyses were made with ImageJ (U.S. NIH, Bethesda, MD; http://imagej.nih.gov/ij/). Results were normalized to β-actin and expressed as arbitrary units.

### ChIP-sequencing

#### FVB Hi-MYC model

ChIP-sequencing was performed as described in Labbé and Zadra *et al*. ^17^. Briefly, fresh-frozen VP tissues from 12-week-old mice were pulverized (Cryoprep Impactor, Covaris), resuspended in PBS + 1% formaldehyde, and incubated at room temperature for 20 minutes. Fixation was stopped by the addition of 0.125 M glycine (final concentration) for 15 minutes at room temperature, then washed with ice cold PBS + EDTA-free protease inhibitor cocktail (PIC; #04693132001, Roche). Multiple biological replicates were combined for each condition in two distinct pools (replicates). Chromatin was isolated by the addition of lysis buffer (0.1% SDS, 1% Triton X-100, 10 mM Tris-HCl (pH 7.4), 1 mM EDTA (pH 8.0), 0.1% NaDOC, 0.13 M NaCl, 1X PIC) + sonication buffer (0.25% sarkosyl, 1 mM DTT) to the samples, which were maintained on ice for 30 minutes. Lysates were sonicated (E210 Focused-ultrasonicator, Covaris) and the DNA was sheared to an average length of ∼200-500 bp. Genomic DNA (input) was isolated by treating sheared chromatin samples with RNase (30 minutes at 37°C), proteinase K (30 minutes at 55°C), de-crosslinking buffer (1% SDS, 100 mM NaHCO3 (final concentration), 6-16 hours at 65°C), followed by purification (#28008, Qiagen). DNA was quantified on a NanoDrop spectrophotometer, using the Quant-iT High-Sensitivity dsDNA Assay Kit (#Q33120, Thermo Fisher Scientific). On ice, AR (2 μg, #ab108341, Abcam), FOXA1 (6 μg, #ab23738, Abcam), RNA Pol II (4 μg, #sc899, Santa Cruz Biotechnology) or H3K27ac (10 μl, #ab4729, Abcam) antibodies were conjugated to a mix of washed Dynabeads protein A and G (Thermo Fisher Scientific), and incubated on a rotator (overnight at 4°C) with 5 μg (AR, FOXA1, RNA Pol II) or 1.5 μg (H3K27ac) of chromatin. ChIP’ed complexes were washed, sequentially treated with RNase (30 minutes at 37°C), proteinase K (30 minutes at 55°C), de-crosslinking buffer (1% SDS, 100 mM NaHCO3 (final concentration), 6-16 hours at 65°C), and purified (#28008, Qiagen). The concentration and size distribution of the immunoprecipitated DNA was measured using the Bioanalyzer High Sensitivity DNA kit (#5067-4626, Agilent). Dana-Farber Cancer Institute Molecular Biology Core Facilities prepared libraries from 2 ng of DNA, using the ThruPLEX DNA-seq kit (#R400427, Rubicon Genomics), according to the manufacturer’s protocol; submitted the finished libraries to quality control analyses as described in the bulk RNA-seq Methods section; ChIP-seq libraries were uniquely indexed in equimolar ratios, and sequenced to a target depth of 40M reads on an Illumina NextSeq500 run, with single-end 75bp reads.

### mCRPC LuCaP PDXs

ChIP-sequencing for AR (N-20; #sc-816, Santa Cruz Biotechnology), FOXA1 (#ab23738, Abcam) and H3K27ac (#C15410196, Diagenode), was performed at the Dana-Farber Cancer Institute using the protocol described previously ^32,63^.

#### LNCaP MYC model

Published gene expression data (GSE73995; ^15^) was downloaded and reanalyzed.

### Bioinformatics analyses – ChIP-seq

#### Peak calling and data analysis

All samples were processed through the computational pipeline developed at the Dana-Farber Cancer Institute Center for Functional Cancer Epigenetics (CFCE) using primarily open source programs. Sequence tags were aligned with Burrows-Wheeler Aligner (BWA) to build mm9 or hg19 and uniquely mapped, non-redundant reads were retained ^64^. These reads were used to generate binding sites with Model-Based Analysis of ChIP-seq 2 (MACS v2.1.1.20160309), with a q-value (FDR) threshold of 0.01 ^65^. We evaluated multiple quality control criteria based on alignment information and peak quality: (i) sequence quality score; (ii) uniquely mappable reads (reads that can only map to one location in the genome); (iii) uniquely mappable locations (locations that can only be mapped by at least one read); (iv) peak overlap with Velcro regions, a comprehensive set of locations – also called consensus signal artifact regions – in the genome that have anomalous, unstructured high signal or read counts in next-generation sequencing experiments independent of cell line and of type of experiment; (v) number of total peaks (the minimum required was 1,000); (vi) high-confidence peaks (the number of peaks that are tenfold enriched over background); (vii) percentage overlap with known DHS sites derived from the ENCODE Project (the minimum required to meet the threshold was 80%); and (viii) peak conservation (a measure of sequence similarity across species based on the hypothesis that conserved sequences are more likely to be functional). Typically, if a sample fails one of these criteria, it will fail many (locations with low mappability will likely have low peak numbers, many of which will likely be in high-mappability regions, etc.).

#### DNA binding motif analyses

Peaks from each group were used for motif analysis by the motif search findMotifsGenome.pl in HOMER (v3.0.0)31, with cutoff q-value ≤ 1e-10.

#### Sample-sample correlation and differential peaks analysis

Sample-sample correlation and differential peaks analysis was performed by the CoBRA pipeline ^66^. Peaks from all samples were merged to create a union set of sites for each transcription factor and histone mark. Read densities were calculated for each peak for each sample, which were used for comparison of cistromes across samples. Sample similarity was determined by hierarchical clustering using the Spearman correlation between samples. Tissue-specific peaks were identified by DEseq2 with adjusted P ≤ 0.05. Total number of reads in each sample was applied to size factor in DEseq2, which can normalize the sequencing depth between samples.

#### ChIP-seq profiles

Given varying alignment of reads or fragments across samples, coverage track bigwig files were calculated for each sample that reflected the coverage signal and sequencing depth using the Chilin pipeline ^67^. The deepTools v.2.3.5 package computeMatrix further computed the average score for each of the samples. Finally, a profile heat map was created based on the scores at genomic positions within 2 kb upstream and downstream of the AR binding sites. All samples were ranked by the average score. ChIP-seq enrichment for transcription factors and histone marks at the loci of selected genes were visualized and plotted using karyoploteR (1.12.4) R package ^68^.

#### RNA Pol II analysis

RNA Pol II traveling ratio (TR) scores for each gene was calculated by comparing the ratio between RNA Pol II density in the promoter region and in the gene region ^41^. The promoter region was defined as -30 bp to +300 bp relative to the transcriptional start site (TSS) and the gene body as the remaining length of the gene. We calculated the bins per million mapped reads (BPM) use bamCoverage and computeMatrix in deepTools v.2.3.5 for promoter and gene body regions. The TR difference between WT and MYC were calculated by TR value in WT minus TR value in MYC. Ranking plot of the WT MYC TR difference for all Pol II bound genes revealed a clear point in the distribution of travel ratio difference where the difference began increasing/decreasing rapidly. To geometrically define this point, we found the x-axis point for which a line with a slope of 1 was tangent to the curve. We defined 246 genes above the increasing point to be pause release genes and 556 genes below the decreasing point to be the pause genes by MYC overexpression. DeepTools function plotProfile and plotHeatmap were used to create the Pol II occupancy (the region ± 3 kb from the start and end of the gene) summary profiles and heatmaps. Kolmogorov-Smirnov test is applied to the TR distribution difference between WT and MYC for Hallmark Androgen_response genes.

### Epigenomics and transcriptomics integration

All genes within the 100 kb of gained AR binding sites in MYC samples were selected, k-means clustering of 3 was applied. Cells were ordered by the pseudotime. GSEA analysis was done using the gene sets deposited in the GSEA website (https://www.gsea-msigdb.org/gsea/msigdb/annotate.jsp). Binding and expression target analysis (BETA) was used to integrates ChIP-seq of transcription factors with differential gene expression data and infer the dysregulated genes ^39^.

### Prostate cancer clinical datasets analyses

#### The Cancer Genome Atlas (TCGA)

RNA-seq readcount and clinical data from 488 samples with prostate cancer (PRAD) were downloaded from the cancer Genome Atlas (TCGA) database (https://cancergenome.nih.gov/) using Bioconductor package TCGAbiolinks (2.14.1)^69^. To calculate transcriptional signature scores, RNA-seq data was normalized to sequencing depth and TPM transformed. Hallmark Androgen_response and Hallmark MYC_targets_V1 gene sets were downloaded from MSigDB ^70^. The AR-A signature comprising nine canonical AR transcriptional targets (*KLK3, KLK2, FKBP5, STEAP1, STEAP2, PPAP2A, RAB3B, ACSL3, NKX3-1*) was derived from previous published work ^34^. Transcriptional signature scores were computed for every patient based on a non-parametric, rank-based method implemented in singscore (1.6.0) R package ^71^. TCGA patients were assigned to the low or high group according to the cut-off point estimated by maximally selected rank statistic maxstat (0.7-25) R package of each signature ^72^. Survival analysis was conducted using survival (3.2-3) R package ^73^, Kaplan-Meier were plotted using survminer (0.4.8) R package ^74^ and log-rank test was used to evaluate the overall statistical significance as well as the comparison between groups. Benjamini-Hochberg was used to correct for multiple testing.

#### Validation cohort

The META855 cohort containing 855 patients treated with radical prostatectomy with available transcriptomic, clinicopathological, and outcomes data selected from five published studies of the Decipher prostate genomic classifier test as previously described ^35^. Microarray expression levels were normalized using the SCAN algorithm (SCAN.UPC package)^75^. The combination of the Hallmark Androgen_response / Hallmark MYC_targets_V1 and AR-A / Hallmark MYC_targets_V1 signatures and their association with BCR and metastatic progression was examined in the META855 cohort using the thresholds obtained from quantiles defined in the TCGA dataset. Patients were divided in four groups and Kaplan-Meier analysis and log-rank test were conducted to evaluate differences in biochemical recurrence and metastatic progression. The prognostic association between the signatures and the clinicopathological factors was assessed using Cox proportional hazard modeling.

#### Castration-resistant prostate cancer

Published gene expression data (GSE126078; ^37^) was downloaded and data analysis was performed using VIPER ^57^.

## Data availability

Data are available from the corresponding authors upon request. The murine (bulk RNA-seq, scRNA-seq and ChIP-seq) and LuCaP PDXs (ChIP-seq) sequencing data reported in this paper will be deposited on NCBI Gene Expression Omnibus (GEO).

## Graphical summary

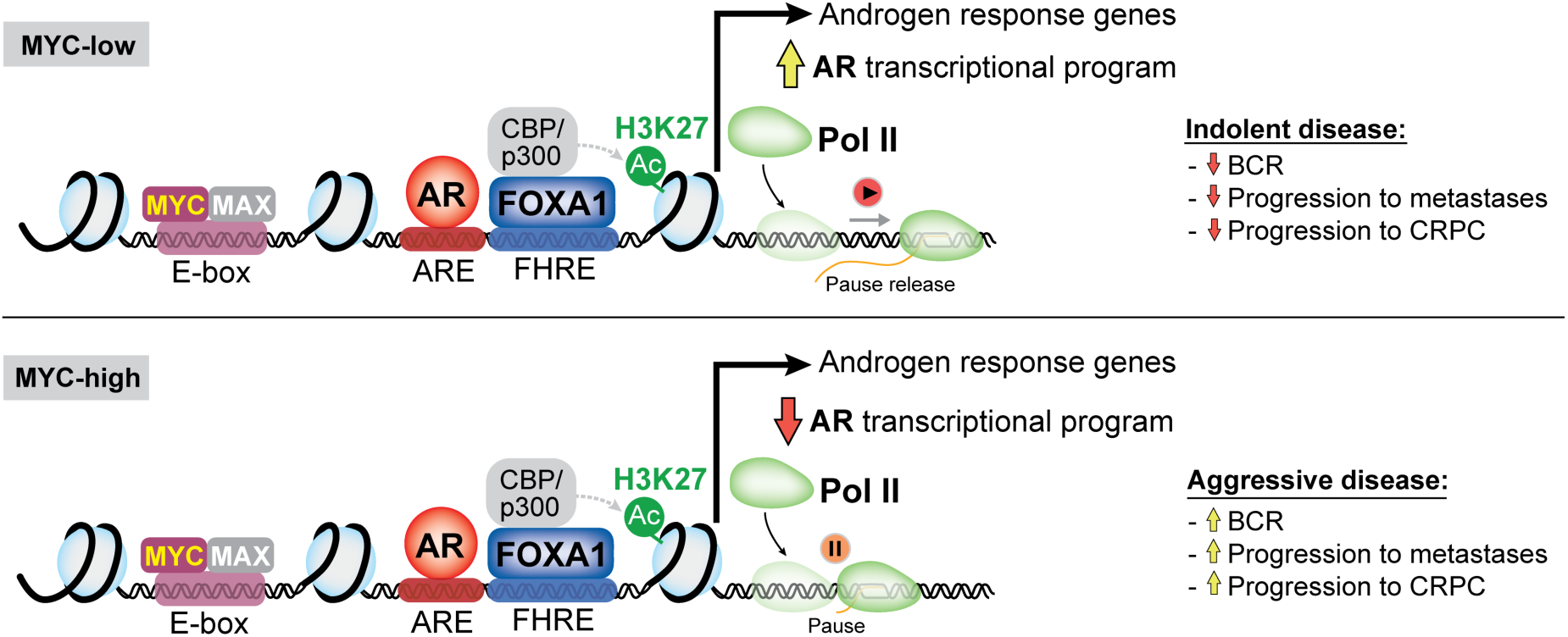

## SUPPLEMENTARY INFORMATION

### SUPPLEMENTARY FIGURES

**Figure S1:**
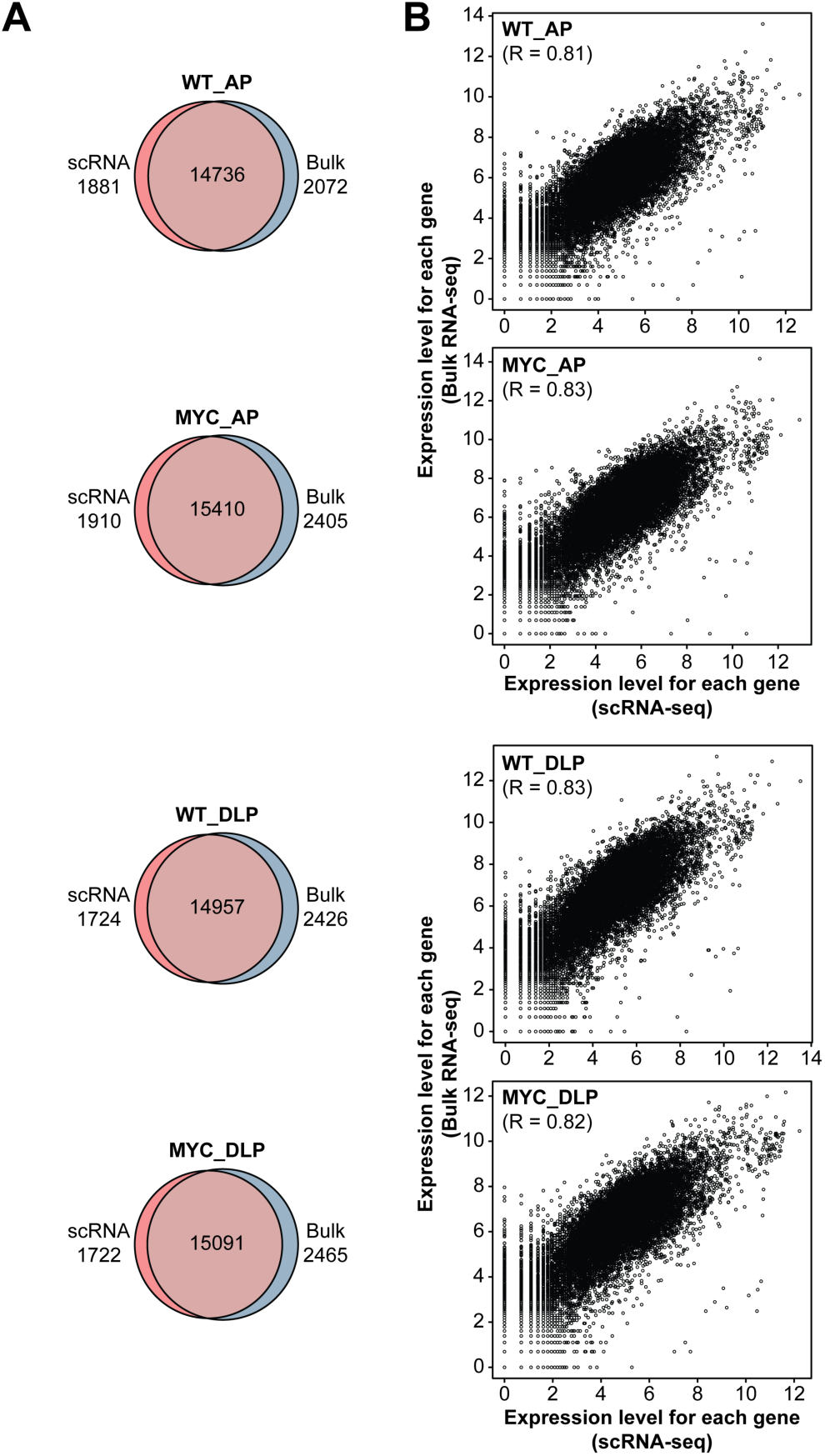
Single-cell transcriptome is highly correlated with bulk gene expression in AP and DLP lobes. (**A, B**) Transcriptional profiling of WT and MYC-transformed AP and DLP lobes reveals high concordance for the total number of genes detected (**A**) and their expression levels (**B**) between bulk and single-cell RNA-seq. WT: wild-type; AP: anterior prostate; DLP: dorsolateral prostate.

**Figure S2:**
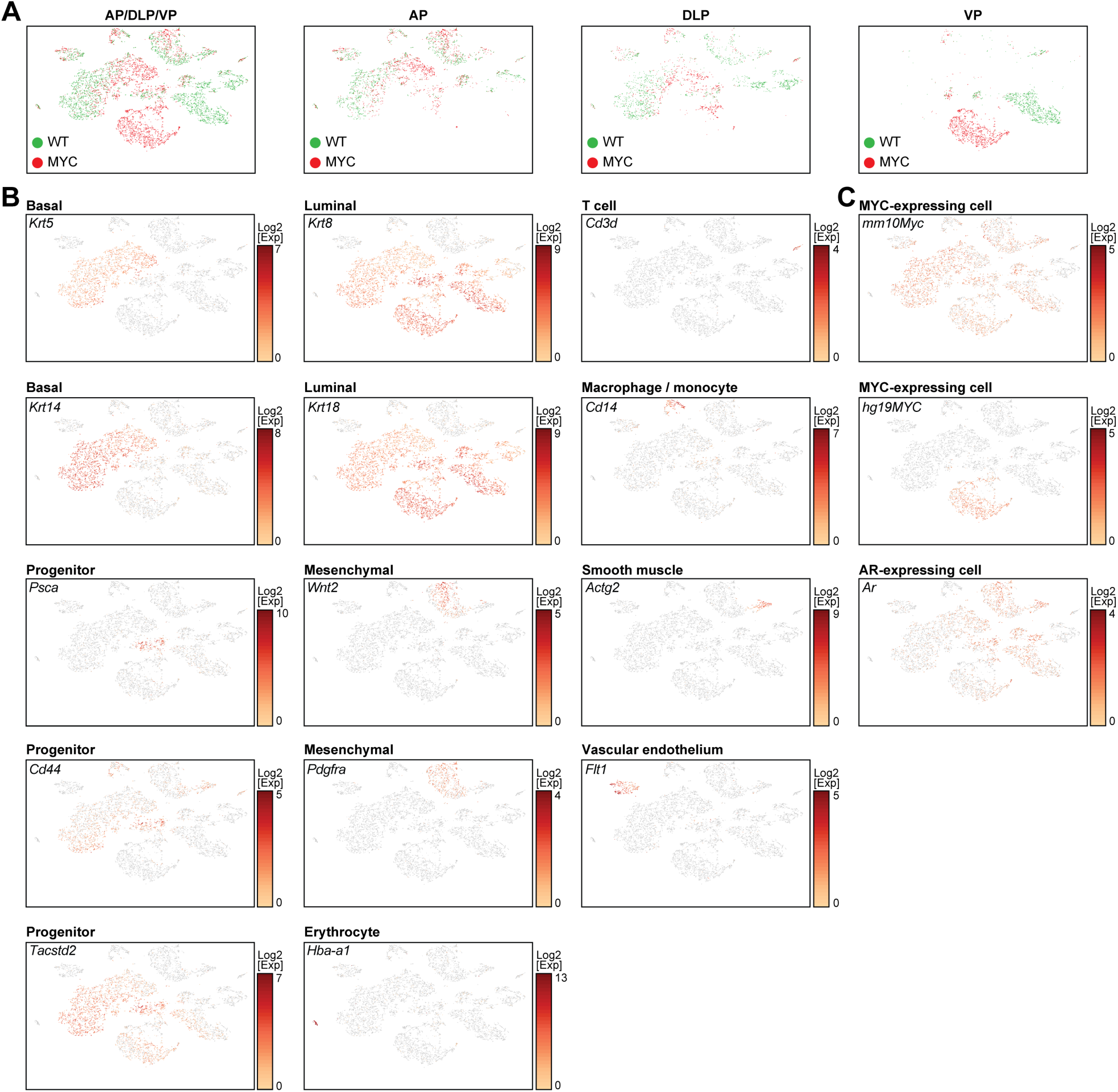
Molecular characterization of murine WT and MYC-transformed prostate lobes. **(A)** Single-cell census of WT and MYC-transformed AP, DLP and VP. tSNE of scRNA-seq profiles (as in Figure 2A), colored by genotype. (**B, C**) Expression of selected markers of different subsets (**B**) as well as murine *Myc* (*mm10Myc*), human *MYC* (*hg19MYC*) and the *Ar* (**C**). WT: wild-type; VP: ventral prostate; DLP: dorsolateral prostate; AP: anterior prostate.

**Figure S3:**
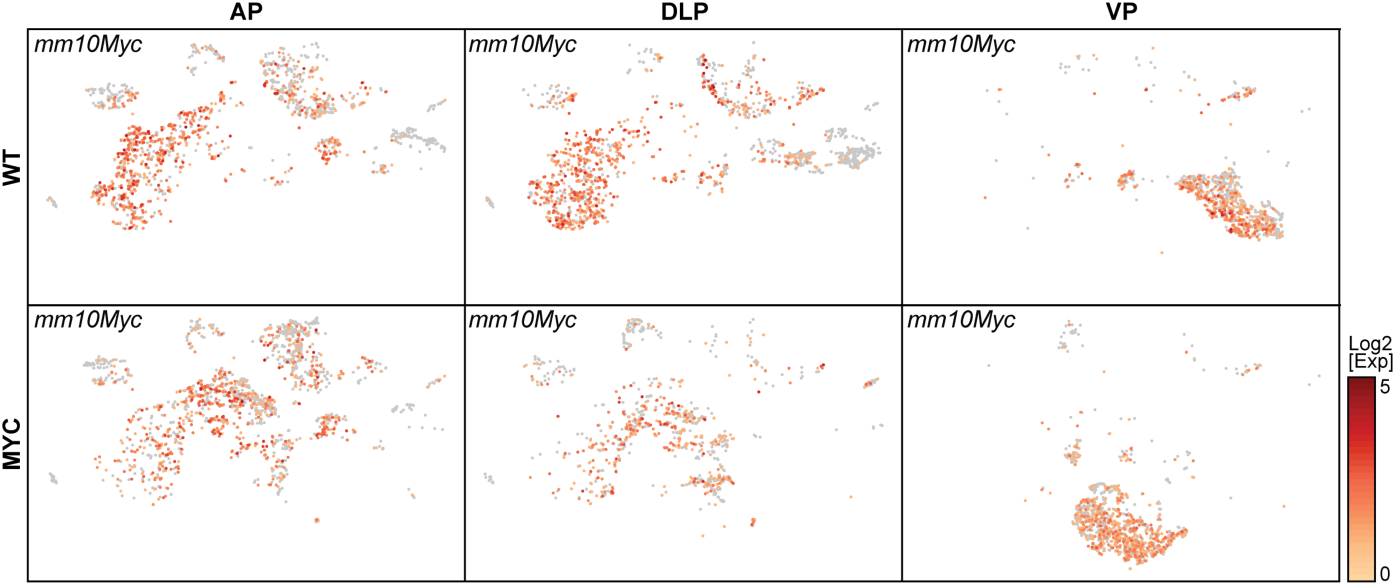
Murine *Myc* is expressed across cell subpopulations and prostate lobes. Expression of murine *Myc* (*mm10Myc*) in WT and MYC-transformed AP, DLP and VP. WT: wild-type; AP: anterior prostate; DLP: dorsolateral prostate; VP: ventral prostate.

**Figure S4:**
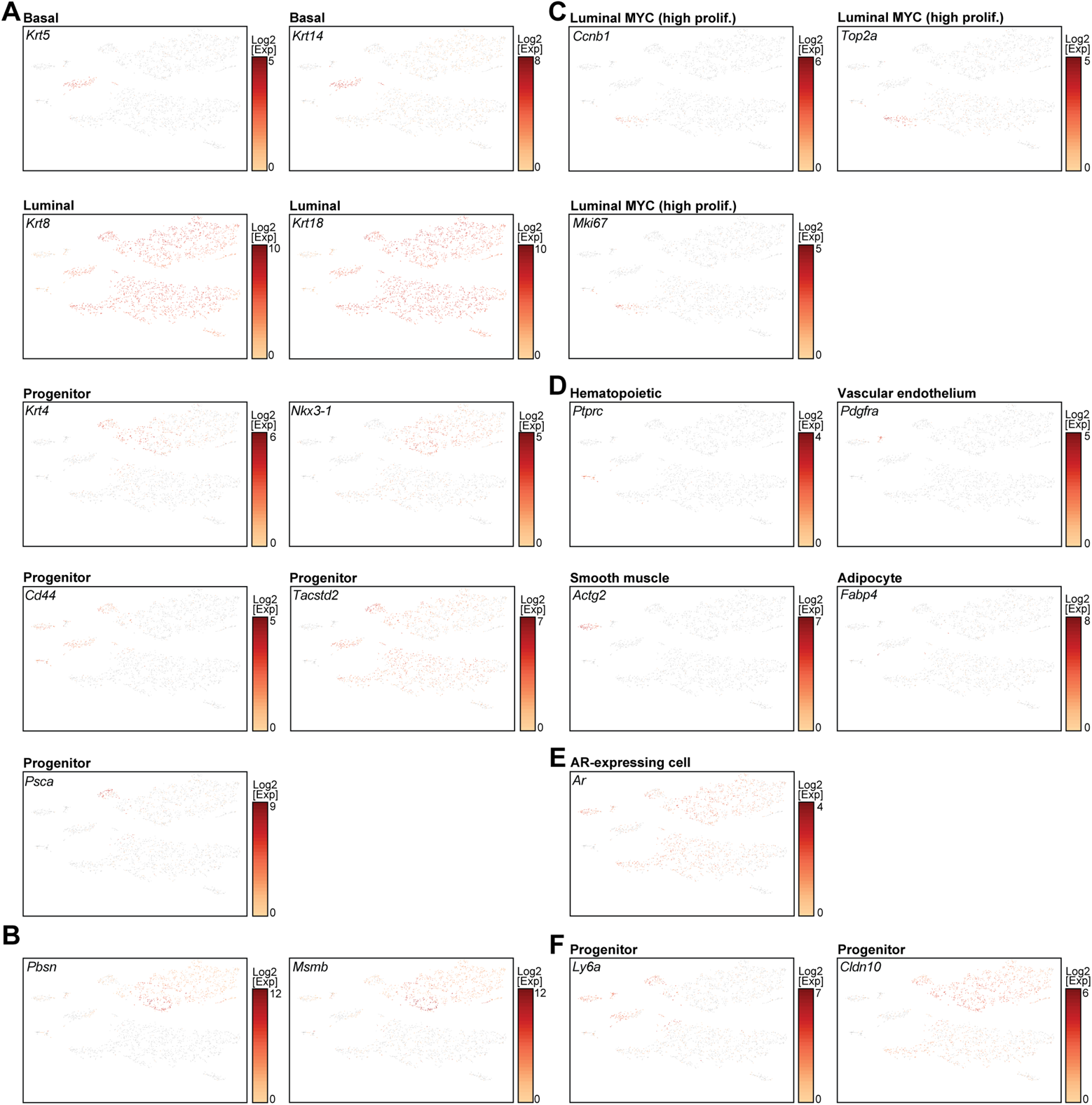
Molecular characterization of murine WT and MYC-transformed VP. **(A-F)** Expression of selected markers of different subsets. WT: wild-type; VP: ventral prostate.

**Figure S5:**
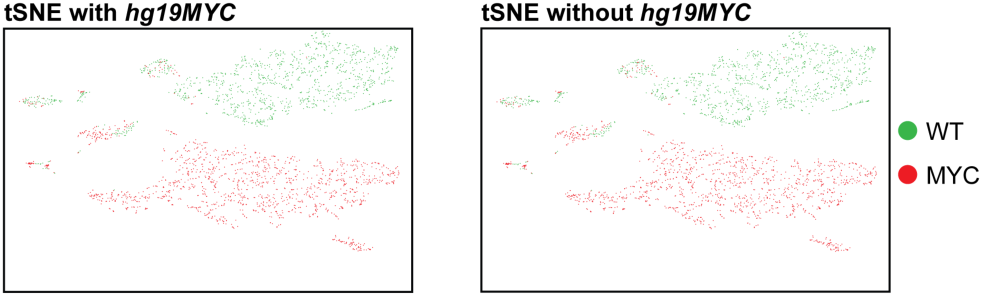
tSNE of scRNA-seq profiles is not affected by the inclusion of human *MYC* transcript. tSNE of VP generated with (*left*) or without (*right*) the inclusion of human *MYC* (*hg19MYC*). WT: wild-type; VP: ventral prostate.

**Figure S6:**
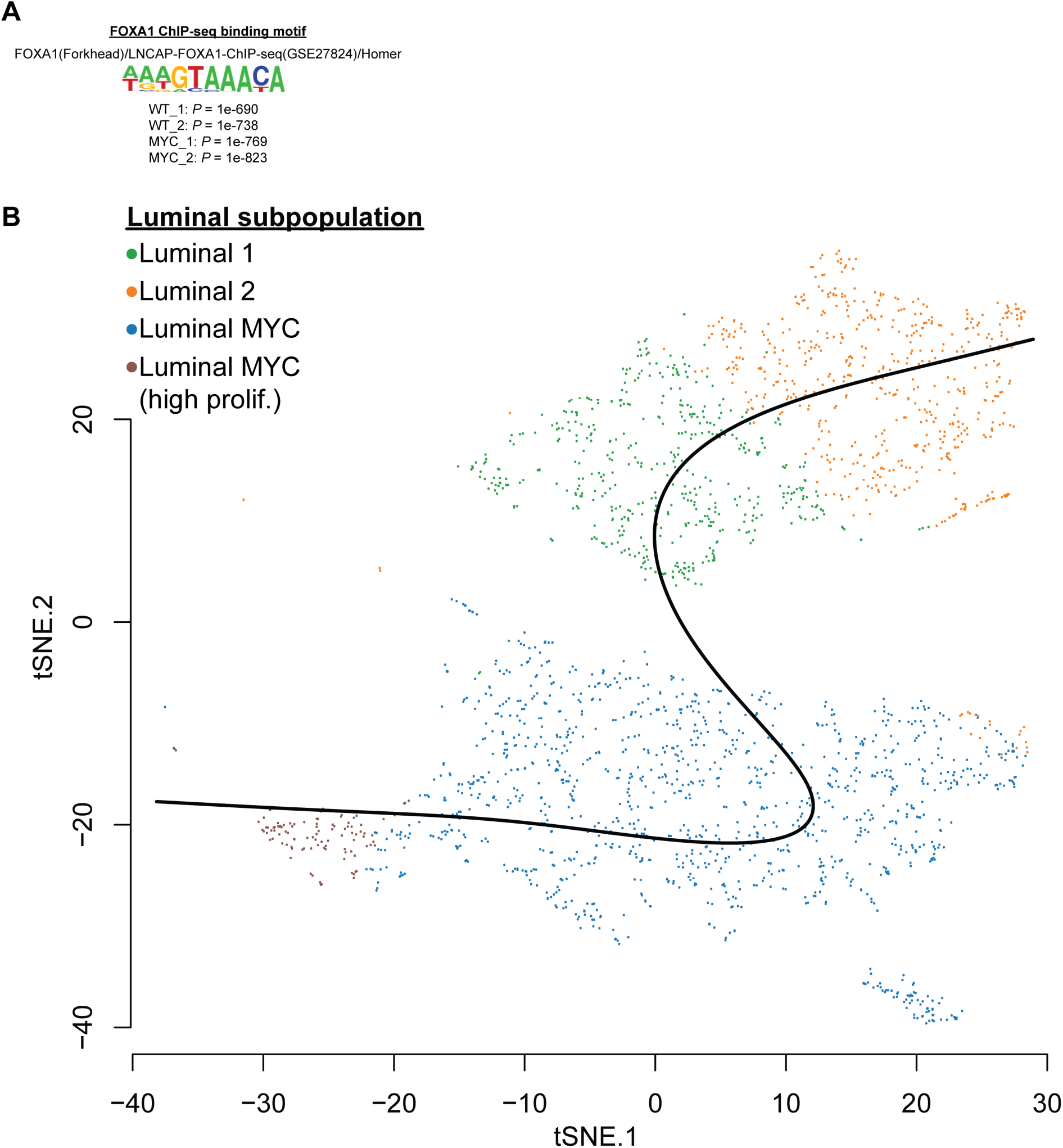
Integration of ChIP-seq with scRNA-seq. (**A**) FOXA1 ChIP-seq identifies FHRE as the top FOXA1 binding motif in WT and MYC-transformed VP. (**B**) Slingshot pseudotime inference used to order luminal cells in **Figure 4G**. WT: wild-type; VP: ventral prostate; FHRE: forkhead response element.

**Figure S7:**
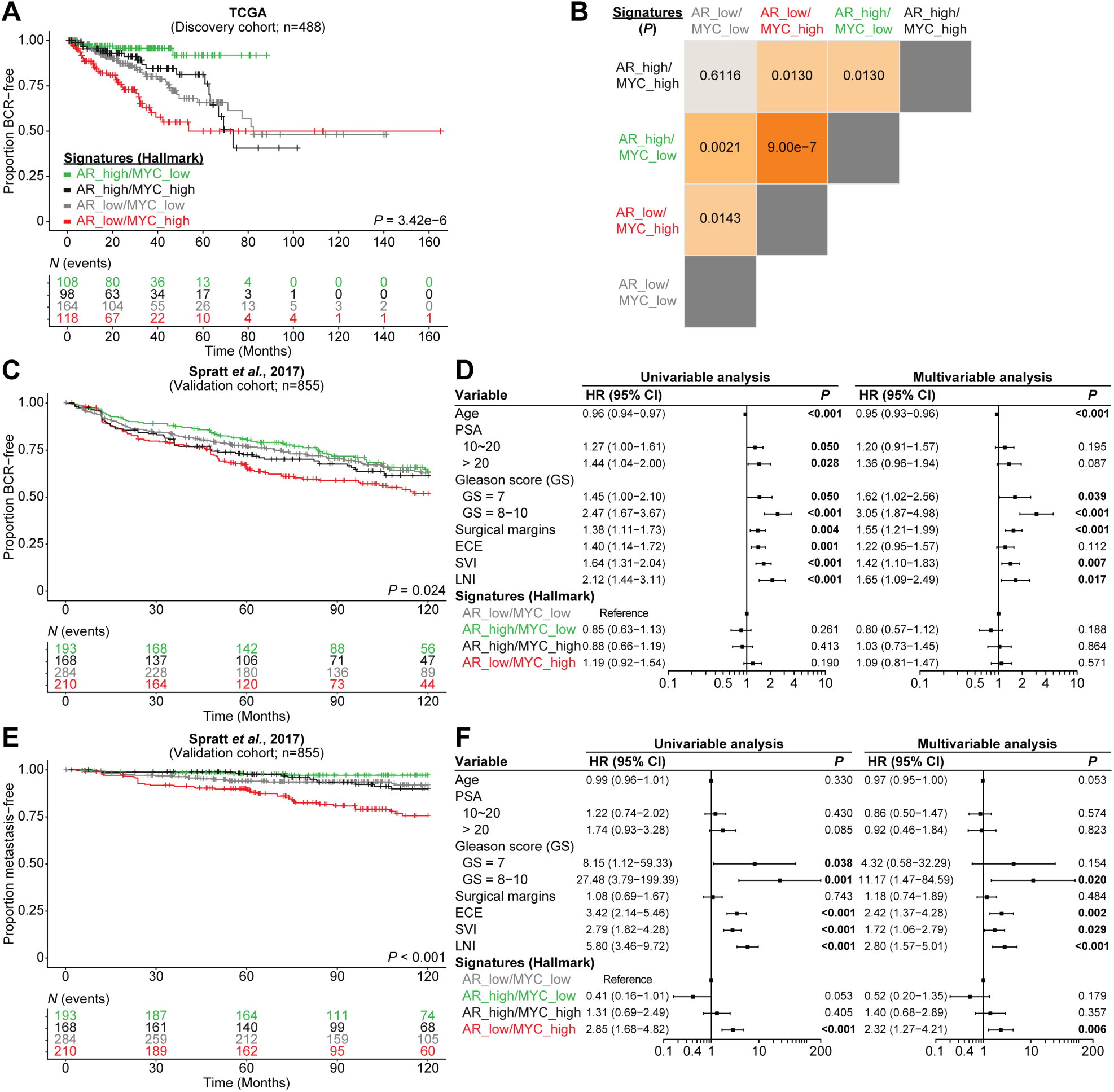
Divergent MYC and AR transcriptional programs dictate disease progression. (**A, B**) Kaplan-Meier curves (**A**) and log-rank tests (**B**) reveal that patients bearing a primary tumor characterized by low AR transcriptional signature (Hallmark) and concurrent high MYC transcriptional signature (Hallmark) have a shorter time to biochemical recurrence (BCR) within the discovery cohort (TCGA). (**C, D**) Kaplan-Meier curves (**C**) but not univariable and multivariable analysis (**D**) confirms that tumors with concurrent low AR and high MYC transcriptional signatures have a significant more rapid development of BCR than tumors with low AR transcriptional signature without an active MYC transcriptional program in the validation cohort (Spratt *et al*., 2017). (**E, F**) Kaplan-Meier curves (**E**), univariable and multivariable analyses (**F**) reveal that tumors with concurrent low AR and high MYC transcriptional signatures are more likely to develop metastatic disease. PSA: prostate-specific antigen; HR: hazard ratio; GS: Gleason score; ECE: extracapsular extension; SVI: seminal vesicles invasion; LNI: lymph node involvement.

**Figure S8:**
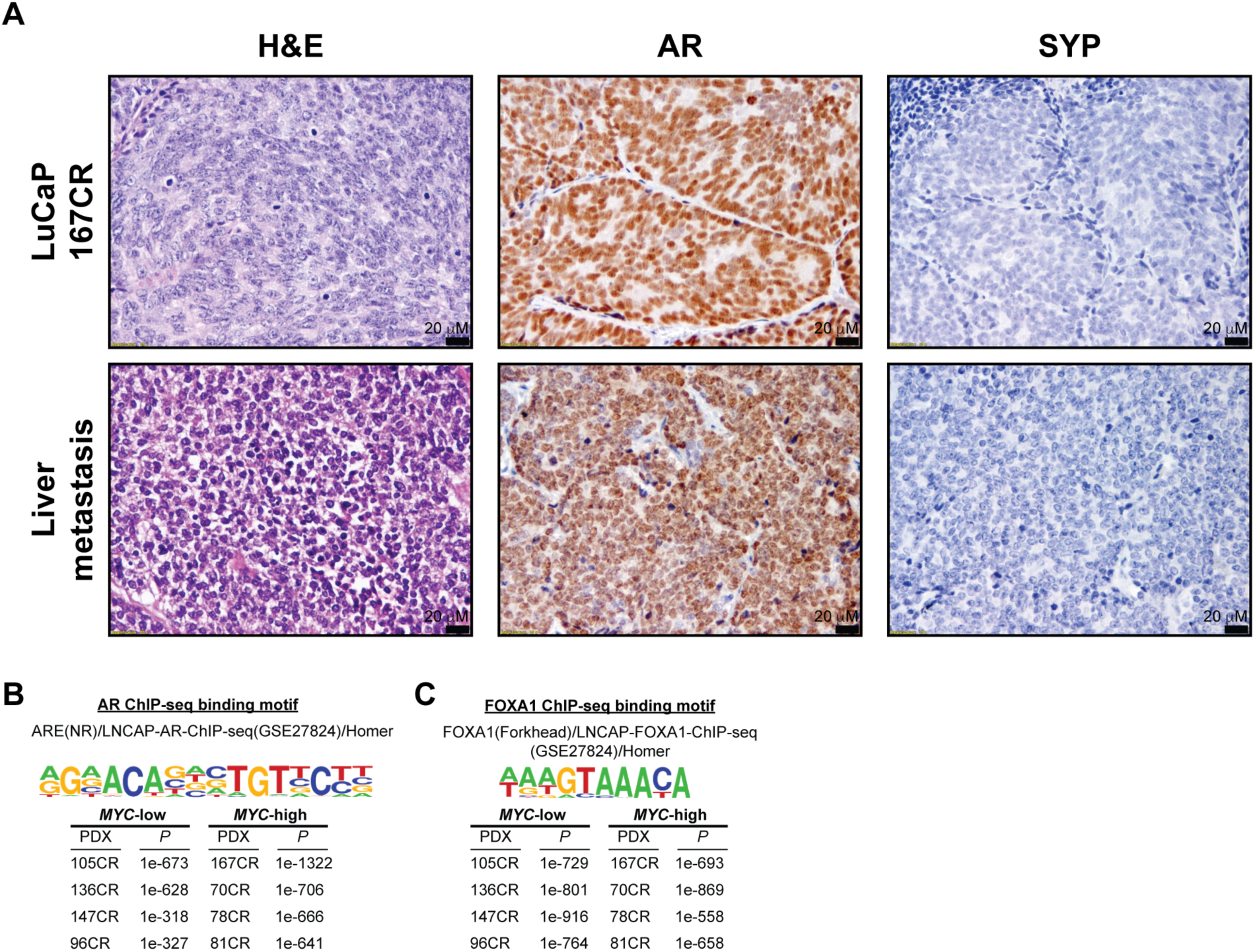
mCRPC LuCaP patient-derived xenograft (PDX) 167CR characterization and top binding motifs in LuCaP PDXs AR and FOXA1 ChIP-seq. (**A**) LuCaP PDX 167CR was established from a liver metastasis of 77-year old Caucasian male who died of abiraterone-, carboplatin- and docetaxel-resistant CRPC. LuCaP 167CR expresses AR, responds to castration and is negative for synaptophysin. Morphology of the PDX was concordant with the original liver metastasis (**B, C**) AR and FOXA1 ChIP-seq identifies ARE (**B**) and FHRE (**C**), respectively as the top binding motif in mCRPC LuCaP PDXs. ARE: androgen response element; FHRE: forkhead response element.

**Figure S9:**
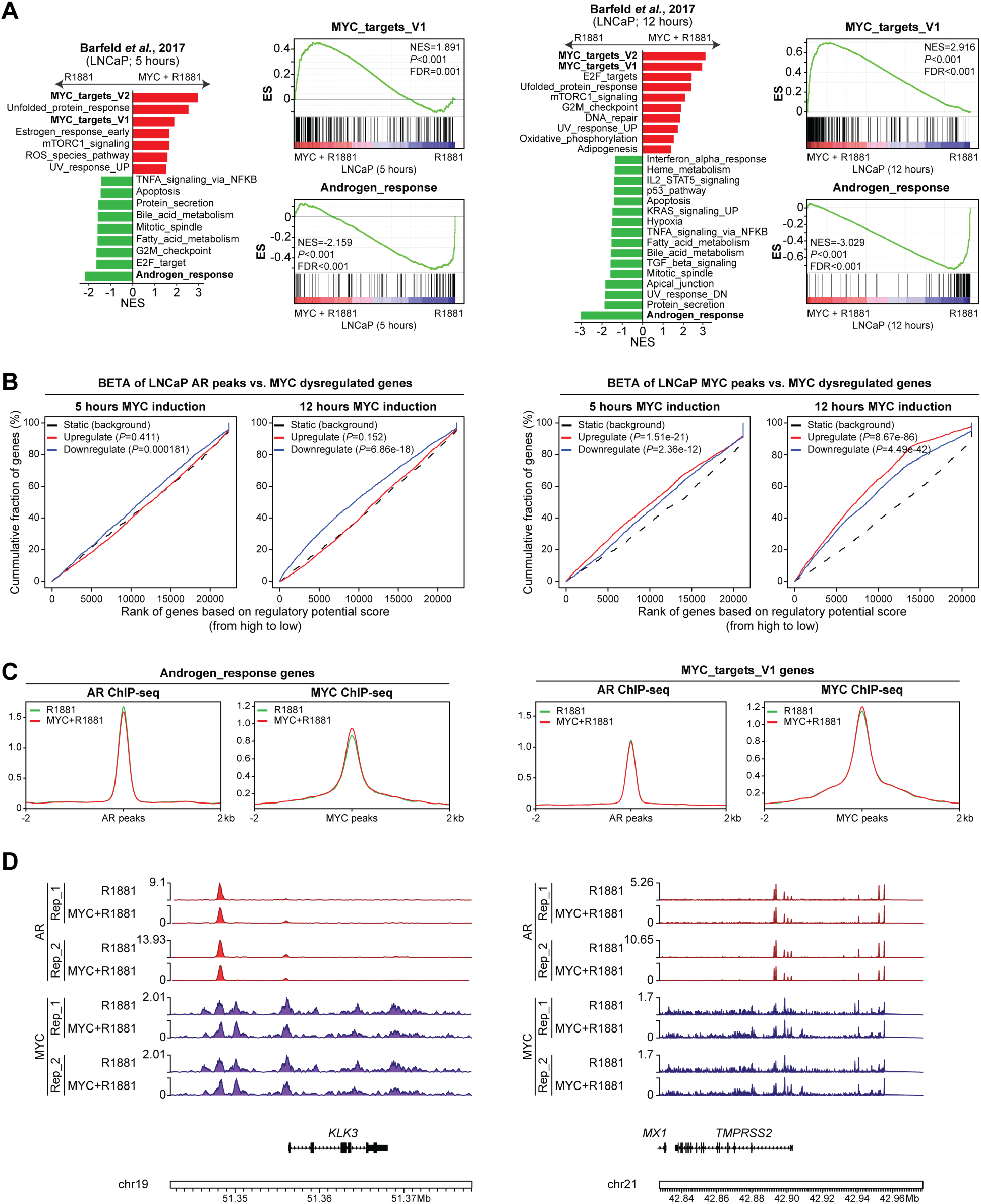
MYC overexpression disrupts the AR transcriptional program in LNCaP cells. Reanalysis of transcriptomics and epigenetics data from Barfed and colleagues ^15^. (**A**) Gene Set Enrichment Analysis (GSEA, Hallmark, P<0.05 and FDR<0.1) revealed an enriched MYC transcriptional program and a depleted AR response following 5 or 12 hours of MYC induction. (**B**) BETA analysis revealed that AR binding sites are associated with gene downregulation while MYC binding sites are associated with gene upregulation following MYC induction. (**C**) AR and MYC binding nearby Androgen_response and MYC_targets_V1 genes is unchanged following MYC induction despite a dampened AR and a heightened MYC transcriptional program. (**D**) Unchanged AR and MYC binding at *KLK3* and *TMPRSS2* loci, AR-dependent genes downregulated by MYC overexpression.

**Figure S10:**
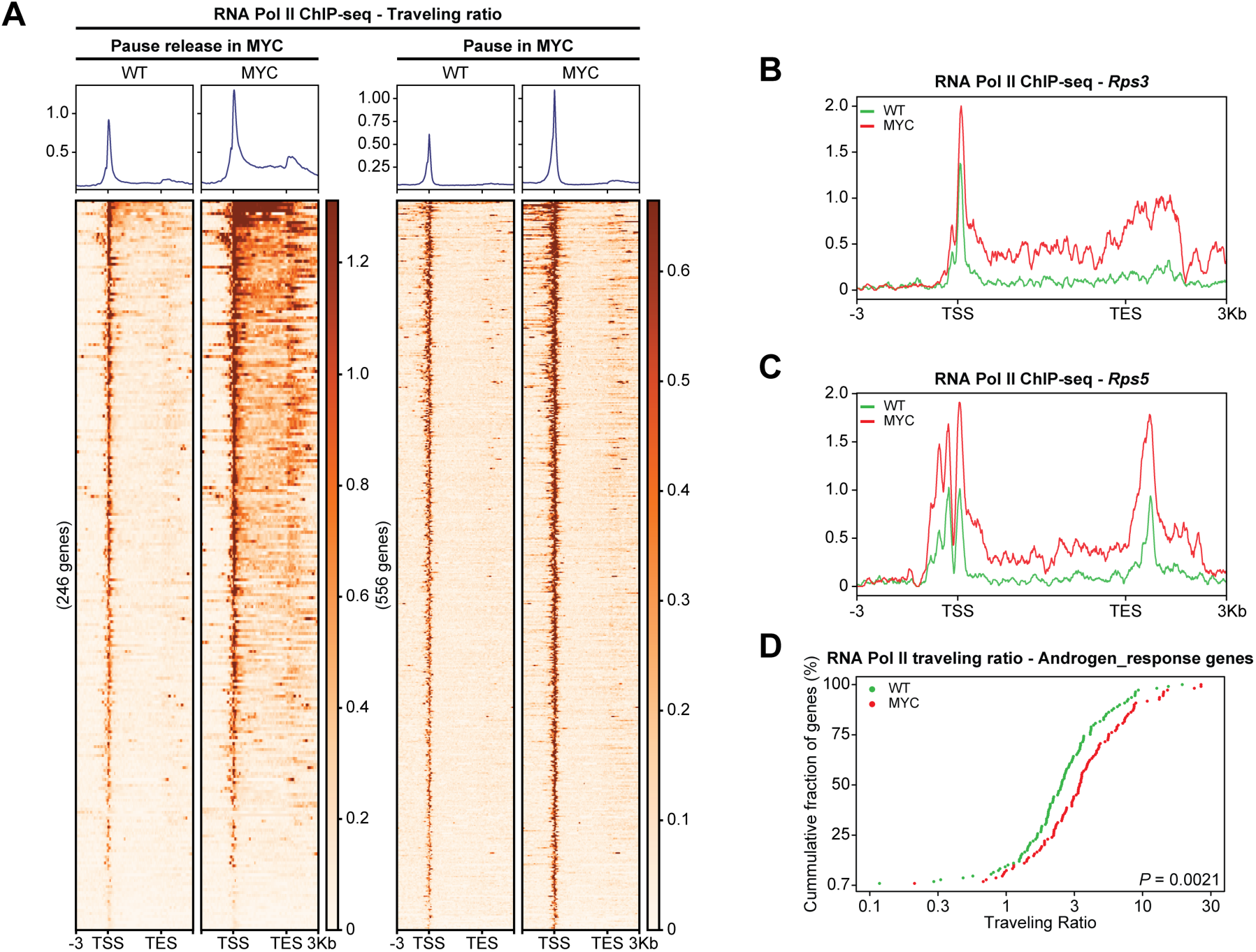
RNA Pol II promoter-proximal pausing. (**A**) RNA Pol II occupancy at pause release (*left*) and pause genes (*right*) following MYC overexpression. (**B, C**) Pause release at *Rps3* (**B**) and *Rps6* (**C**) MYC_targets_V1 genes. (**D**) RNA Pol II traveling ratio reveals greater promoter-proximal pausing at Androgen_response genes (non-smoothed curves).

### SUPPLEMENTARY DATA

**Supplementary Data 1:**
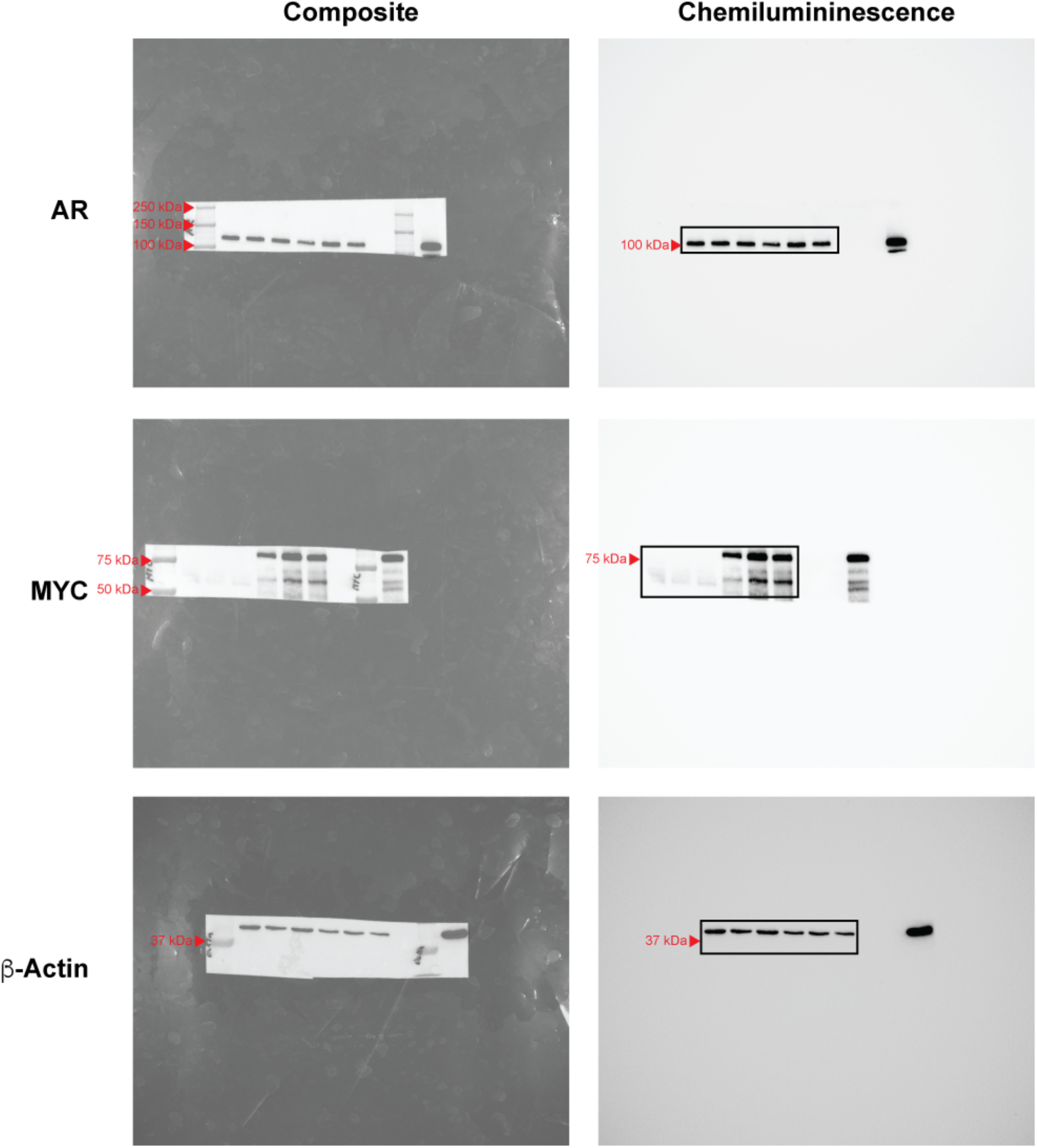
Unprocessed western blots from **Figure 3E**.

**Supplementary Data 2:**
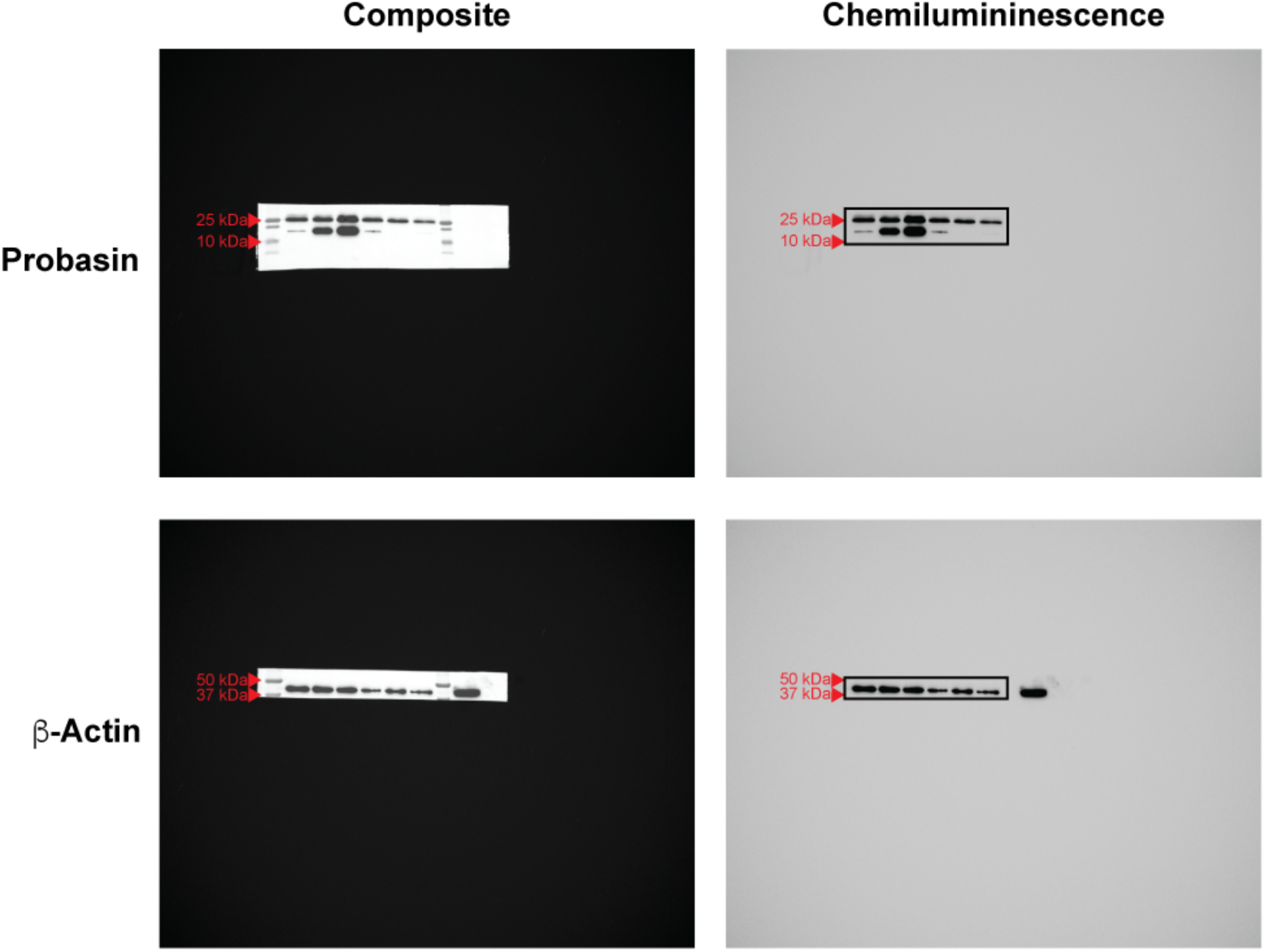
Unprocessed western blots from **Figure 7E**.

